# Multigenerational exposure to heat stress induces phenotypic resilience, and genetic and epigenetic variations in *Arabidopsis thaliana* offspring

**DOI:** 10.1101/2020.11.30.405365

**Authors:** Narendra Singh Yadav, Viktor Titov, Ivie Ayemere, Boseon Byeon, Yaroslav Ilnytskyy, Igor Kovalchuk

## Abstract

Plants are sedentary organisms that constantly sense changes in their environment and react to various environmental cues. On a short-time scale, plants respond through alterations in their physiology, and on a long-time scale, plants alter their development and pass on the memory of stress to the progeny. The latter is controlled genetically and epigenetically and allows the progeny to be primed for future stress encounters, thus increasing the likelihood of survival. The current study intended to explore the effects of multigenerational heat stress in *Arabidopsis thaliana.* 25 generations of *Arabidopsis thaliana* were propagated in the presence of heat stress. The multigenerational stressed lineage F25H exhibited a higher tolerance to heat stress and elevated frequency of homologous recombination, as compared to the parallel control progeny F25C. A comparison of genomic sequences revealed that the F25H lineage had a three-fold higher number of mutations (SNPs and INDELs) as compared control lineages, suggesting that heat stress induced genetic variations in the heat-stressed progeny. The F25H stressed progeny showed a 7-fold higher number of non-synonymous mutations than the F25C line. Methylome analysis revealed that the F25H stressed progeny showed a lower global methylation level in the CHH context than the control progeny. The F25H and F25C lineages were different from the parental control lineage F2C by 66,491 and 80,464 differentially methylated positions (DMPs), respectively. F25H stressed progeny displayed higher frequency of methylation changes in the gene body and lower in the body of transposable elements (TEs). Gene Ontology analysis revealed that CG-DMRs were enriched in processes such as response to abiotic and biotic stimulus, cell organizations and biogenesis, and DNA or RNA metabolism. Hierarchical clustering of these epimutations separated the heat stressed and control parental progenies into distinct groups which revealed the non-random nature of epimutations. We observed an overall higher number of epigenetic variations than genetic variations in all comparison groups, indicating that epigenetic variations are more prevalent than genetic variations. The largest difference in epigenetic and genetic variations was observed between control plants comparison (F25C vs F2C), which clearly indicated that the spontaneous nature of epigenetic variations and heat-inducible nature of genetic variations. Overall, our study showed that progenies derived from multigenerational heat stress displayed a notable adaption in context of phenotypic, genotypic and epigenotypic resilience.

## Introduction

Plants are continuously exposed to their surrounding environment which effects their growth and impacts agricultural yields (Wei et al. 2020). These interactions result in changes in gene expression and physiological and biomolecular responses that alter a plant’s phenotype (Zheng et al. 2013; Wei et al. 2020).

Plants respond to the environment through the adaptation and tolerance that ensure the survival of both, the plant itself and its progeny (Huang et al. 2012). This phenomenon is feasible due to the plasticity of the plant genome and epigenome, and it evolves through the environmentally induced alterations and the development of phenotypic resilience (Nicotra et al. 2010; Zhang et al. 2018; Lind and Spagopoulou 2018; Miryeganeh and Saze 2020). The phenotypic plasticity in response to environmental conditions is coordinated through the stress perception and activation of signalling pathways and responsive genes (Chinnusamy et al. 2004; Zheng et al. 2013). The adaptation to the environment is not restricted to changes in the physiology of the plant, but it also results in changes in the genome and epigenome (Gutzat and Scheid 2012; Zhang et al. 2018; Miryeganeh and Saze 2020).

Environmental stresses have an impact on the directly exposed generation as well as on their offspring *via* parental effects and/ or transgenerational effects (Herman and Sultan 2011). Parental effects, which are also referred as intergenerational effects, are described by phenotypic changes in the immediate progeny of stressed plants. In contrast, transgenerational effects refer exclusively to changes in the phenotype of the progeny that is separated from the stressed parent by at least one generation, thus representing true inheritance of acquired changes. However, many of the described intergenerational effects share their mechanisms with transgenerational effects, and many publications do not distinguish between the two, often referring to both as transgenerational (Perez and Lehner 2019). The RNA interference-related mechanisms can regulate the transgenerational inheritance of a specific chromatin or DNA modifications (Duempelmann et al. 2020). Very recently, Yang et al. (2020) have reported that the segregation of the *MSH1* RNAi transgene produces the heritable non-genetic memory in association with methylome reprogramming. The *msh1* reprogramming is dependent on functional HDA6 and DNA-methyltransferase MET1, and the transition to memory requires the RdDM pathway. This system of phenotypic plasticity may play a vital role for plant adaptation to the changing environment (Yang et al. 2020).

While the exact mechanisms of stress adaption and transgenerational stress memory remain unclear; the understanding of these processes has far-reaching implications for plant breeding, genetic engineering, and the development of stress tolerant crops (Bilichak and Kovalchuk 2016; Zhang et al. 2018). A substantial number of studies demonstrate the ability to maintain the memory of stress exposure throughout ontogenesis and transmit this memory to the following generation (Boyko and Kovalchuk 2010; Suter and Widmer 2013; Bilichak and Kovalchuk 2016; Ramírez-Carrasco et al. 2017; Zheng et al. 2017).

The mechanism of stress memory most likely includes genetic and epigenetic changes, with the latter ones being more prevalent. The genetic variations within species are identified as single nucleotide polymorphisms (SNPs) and insertions and deletions (INDELs). It has been reported that stress induced the accumulation of genetic mutations in stressed lineages of bacteria (Al Mamun et al. 2012), yeast (Shor et al. 2013) and Arabidopsis (Jiang et al. 2014; Belfield et al. 2020). SNPs may cause the phenotypic diversity such as flowering time adaptation, the colour of plant fruit and tolerance to various biotic and abiotic factors (Vidal et al. 2012; Jang et al. 2015).

In the case of epigenetics, phenotype traits are transferred to the progeny without changes in gene sequence (Eichten et al. 2014; Bilichak and Kovalchuk 2016; Zheng et al. 2017; Lind and Spagopoulou 2018; Miryeganeh and Saze 2020). Epigenetic regulation consists of three primary mechanisms - DNA methylation, histone modifications and small RNAs (sRNAs) activity. DNA methylation is the most established process that alters gene expression utilized by plants for changing their phenotypes in stressful environment, acting as a complementary mode of transferring heritable information (Mirouze and Paszkowski 2011; Schmitz et al. 2011). Many recent reports have demonstrated that plant DNA methylation can be altered at individual loci or across the entire genome under environmental stress conditions (Zhang et al. 2018 and reference therein). In rice, the naturally occurring variations in DNA methylation are well studied epimutations at single gene loci that have been shown to result in heritable morphological variations without altering the DNA sequence of rice genes (Zheng et al. 2017; Miura et al. 2009). The chilling-induced loss of tomato flavor is associated with changes in DNA methylation, including the *RIN* gene (Zhang et al. 2016). In *Arabidopsis*, quantitative resistance to clubroot infection is mediated by transgenerational epigenetic variations (Liégard et al. 2019).

Changes in DNA methylation include differentially methylated positions (DMPs), where single cytosines are involved, and differentially methylated regions (DMRs), where multiple cytosines in a given region (arbitrarily defined) are considered. DMPs and DMRs are specific epimutations that can be associated with the epigenetic inheritance of stress response (Zhang et al. 2018). Exposure to environmental factors can induce changes in the specific DMRs, which suggest the existence of the exposure specific DMRs (Haque et al. 2016; Manikkam et al. 2012). Some examples include DMRs associated with the differentially expressed genes in the regulation of the *Arabidopsis thaliana* immune system against *Pseudomonas syringae* (Dowen et al. 2012) and the location-specific DMRs at specific *cis*-regulatory sites in the *Hevea brasiliensis* tree crop in response to cold and drought stresses (Uthup et al. 2011).

Previous research from our laboratory demonstrated that the progeny of plants exposed to various abiotic and biotic stressors exhibited changes in phenotypic traits such as leaf size and number, flowering time, and seed size (Boyko et al. 2010; Bilichak et al. 2012; Migicovsky et al. 2014). The progeny of stressed plants also displayed a certain degree of stress tolerance and cross-tolerance lasting for one or two generations without the maintenance of stress factor (Rahavi and Kovalchuk 2013; Migicovsky et al. 2014). Our previous studies also demonstrated changes in the genome – rearrangements at the resistance genes loci in the progeny of plants infected with tobacco mosaic virus (Boyko et al. 2007) and in the epigenome – changes in DNA methylation and histone modifications in the progeny of plants exposed to various stresses such as salt (Bilichak et al. 2012), heat (Migicovsky et al. 2014), cold (Migicovsky and Kovalchuk, 2013), UVC (Migicovsky and Kovalchuk, 2014) and viral infection (Boyko et al. 2007). Although these studies did not explicitly analyse the mechanism of transgenerational inheritance, they proposed the role of epigenetic factors and demonstrated genetic changes relevant to stress exposure (Boyko et al. 2010; Boyko and Kovalchuk 2010; Bilichak et al. 2015).

Based on studies conducted in our lab and several other labs, we hypothesized that multigenerational exposure to heat stress would be better in conveying stress memory. We also hypothesized that the plant population propagated under multigenerational heat stress would show a broader genotypic and epigenotypic diversity compared with the population of plants propagated without stress.

In this study, we have shown the ability of Arabidopsis plants to form heat stress memory using the repeated heat stress exposure over 25 consecutive generations. The F25H generation of stressed progeny was compared with the parental F2C and parallel F25C control progenies for heat stress resilience and the presence of heat stress-induced genetic and epigenetic changes.

## Materials and Methods

### Parental generation and progeny plants

Plant seeds used for the parental generation (P) were obtained from single homozygous *Arabidopsis thaliana* (Columbia ecotype, Col-0) plant transgenic for the *luciferase* (LUC) recombination reporter gene (15*d*8 line) carrying a copy of a direct repeat of the *luciferase* recombinant transgene. All parental plants were grown to flower, and seeds were collected and pooled from approximately 16-20 plants per each experimental line. The subsequent progeny plants were obtained from these seeds; specifically, ∼ 100 seeds were sown, and ∼ 20 randomly selected plants were either exposed to heat or grown under normal conditions, both allowed to set seeds, thus forming the next generation. The same process was repeated consecutively until twenty-five generations representing the progeny plant lines (Figure 1A). The seeds were kept in storage under dry conditions at room temperature.

**Figure 1.**
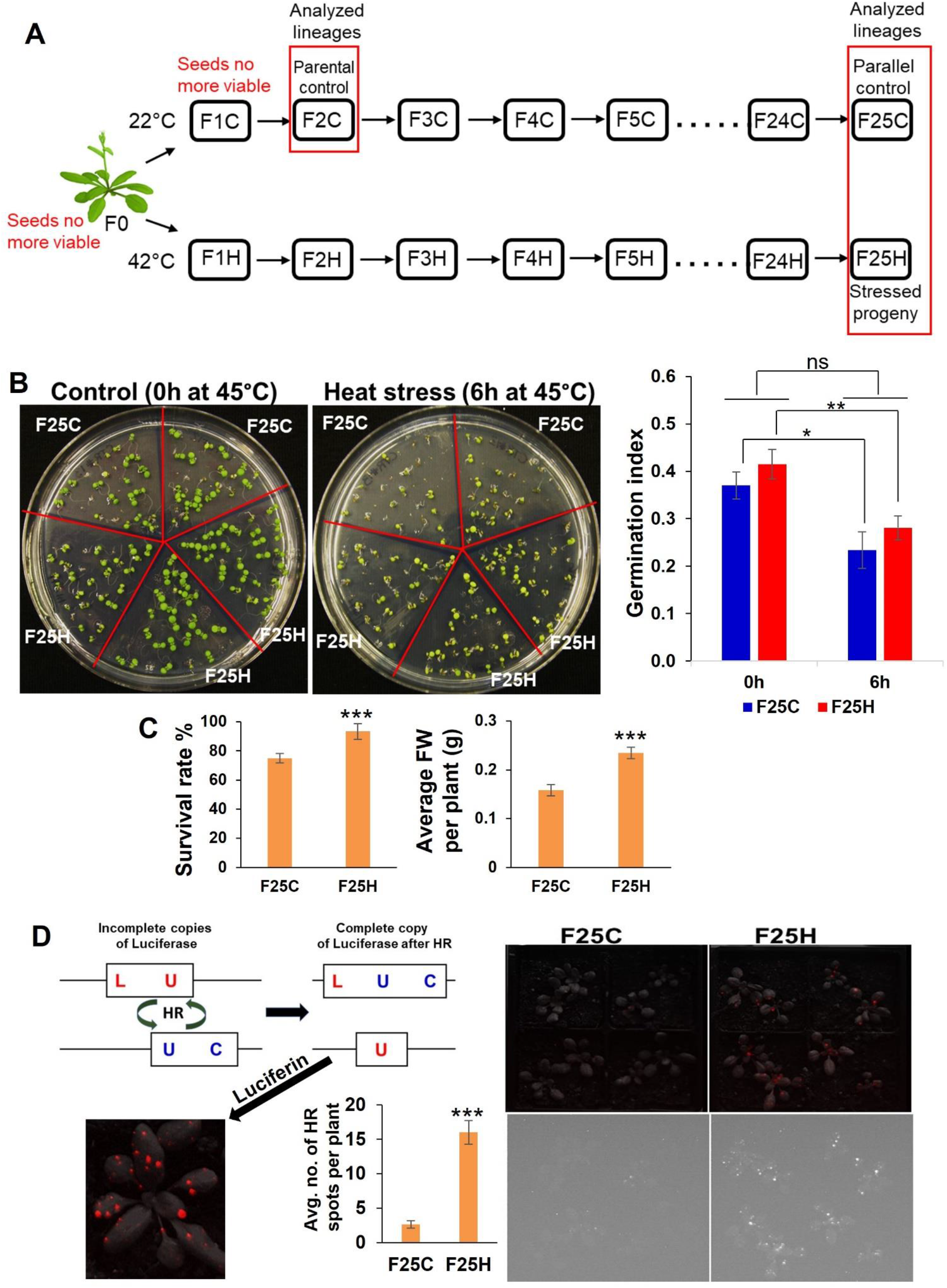
The heat-stressed progeny exhibited an increase in heat stress resilience and higher frequency of recombination events. (**A**) Schematic diagram for propagations of 25 generations of heat stressed and parallel control lineages. Single *Arabidopsis thaliana* plant (Col-0, *15d8 line*) was used for seed origin. Plants were grown for twenty-five generations either exposed to heat stress (42°C) or in a control ambient environment (22°C). The ‘H’ group included the heat-stressed progenies, while the ‘C’ group included plants grown under control conditions acting as a parallel control. Seeds of second-generation F2C were used as parental control because seeds of F0 and F1C were no longer viable. Lineages used in this study are shown in the red box. (**B**) Trends in the germination of the F25H progeny under heat stress. Representative photographs of germination under control (0h) and heat-stress (6h) conditions (the left panel). Seeds were either from the parallel control (F25C) or the heat-stressed progeny (F25H). Each plate represents one set, and in total, five sets were used for each treatment. The right panel: The germination index graph. 0h: non-stress control (0 hour at 45°C); 6h: heat-stress (6 hours at 45°C). Heat-stress was applied at 45°C for 6h, just 4-days after stratification at 4°C. The germination index was measured on the 6^th^ day. The data were analysed by two-way ANOVA with the Tukey’s multiple comparisons test using GraphPad Prism version 8.4.2 for Windows. The data are shown as an average mean ±SD. Mean values that were significantly different from each other are indicated by * (*p <0.05*), ** (*p <0.01*) and those that were non-significant are indicated by ns (*p >0.05*). (**C**) Left panel shows survival of mature plants of stressed progeny (F25H vs F25C) under heat-stress. A survival rate was calculated after 10-days of recovery and shown as an average percentage (with ±SD) of plants surviving the stress from three independent experiments. Right panel shows the average fresh weight (FW) per plant (with ±SE) of survived plants. The data were analysed by one-way ANOVA with Tukey’s multiple comparisons test using GraphPad Prism version 8.4.2 for Windows. The asterisk shows a significant difference between the stressed progeny and parallel control progeny (*** *p <0.001*). (**D**) Somatic HRF under non-stress conditions. Progeny of stressed plants exhibited a higher somatic homologous recombination (HRF) without exposure to heat-stress. Left panel shows the scheme of the HRF analysis. Right panel shows the recombination events in F25C and F25H plants. The upper panel is a superimposed photograph of HR events and plants, and the lower panel is an original florescence photograph. Left panel also shows the average no. of HR spots per plant (with ±SE). The data were analysed by one-way ANOVA with Tukey’s multiple comparisons test using GraphPad Prism version 8.4.2 for Windows. The asterisk shows a significant difference between the stressed progeny and parallel control progeny (*** *p <0.001*).

The progeny plants obtained from the progeny seeds through the process described above for every generation had two groups: the stressed (H) and non-stressed groups (C) which acted as the parallel control group within each generation. All progeny plants representing the ‘H’ group were subjected to 42°C heat stress for 4 hours per day for four consecutive days, while the trays with plants were covered with plastic domes to prevent loss of moisture, starting at day 10 post germination, and they were grown to flower and set seeds. Seeds were then collected and propagated to obtain the next successive generation, thus creating 25 generations of the heat stressed ‘H’ group. On the other hand, the progeny plants representing the ‘C’ group were grown under normal conditions and set seeds; this was also done for 25 generations.

### Plant growth conditions

The seeds were kept at 4°C for seven days to initiate stratification, and then sown in all-purpose potting soil prepared with water containing a generic fertilizer (Miracle-Gro, Scotts Canada). The moist soil containing seeds was further stratified for 48 hours at 4°C. Plants were grown in a growth chamber (BioChambers, Manitoba, Canada) at 22°C under the extended day conditions of 16 hours light (120 μmol photons m^−2^ s^−1^) and 8 hours dark. Approximately three to five days post germination (dpg), plant seedlings were transplanted into individual pots containing a 9:1 ratio of soil to vermiculite composition prepared with a fertilizer. Pots containing plants were placed in trays and watered from below.

### Lineages used for heat stress phenotyping

The seeds of F0 (parental) and F1 generations were no longer viable, hence F2C seeds were used as parental control. Overall, the lineages F2C (parental control), F25H (the 25^th^ generation of stressed progeny) and F25C (the 25^th^ generation of parallel control progeny) were used for this study.

### Phenotypic analysis under heat stress treatment

The F25H stressed progenies were studied for heat stress memory and response. The response to heat stress was determined at the germination or mature plant stage. For germination experiment, three replicates were used for the stressed progeny, and two replicates were used for the parallel control progeny in one MS media plate. Each plate represented one set, and in total, five sets were used for each treatment. Heat stress was applied at 45°C for 6 h, just 4 days after stratification at 4°C. The percentage of germination and the speed of germination were qualitatively and quantitatively evaluated between the groups. To analyse the heat stress tolerance at the seedling stage, we exposed F25C and F25H plants to 42°C for 4h for four consecutive days (keeping plants under plastic domes – in the same way as they were propagated under stress for 25 generations). To test the heat stress phenotype at the mature plant stage, the plants were grown under normal condition, and heat stress was applied just before bolting for 96 h at 40°C. The qualitative and quantitative assessment of survived plants has been done after 10-days of recovery from stress. Phenotype changes such as the survival rate and fresh weight of the survived plants were measured.

### The analysis of homologous recombination frequency (HRF)

The *Arabidopsis thaliana #*15*d*8 line transgenic for the *luciferase* (LUC) recombination reporter gene enables the analysis of HRF. Recombination events occur *via* the rearrangement between two homologous inactive copies of the luciferase transgene. The scheme of the HRF analysis is shown in Figure 1D (left panel). The F25H and F25C plants were analysed for HRF under non-stress conditions. HR events are analysed by scoring bright sectors on a dark background with a CCD camera after spraying with luciferin (Ilnytskyy et al. 2004). These sectors represent cells in which recombination events occurred. For the HRF analysis, we used at least 50 three-week-old plants per lineages. Each experiment was repeated at least twice.

### Lineages used for genomic and epigenomic study

The parental F2C plants and the progeny of 25^th^ generation (both stressed progeny F25H and parallel control progeny F25C) plants were grown to about 21 days post germination (dpg), rosette leaves tissues were harvested from individual plants and snap frozen in liquid nitrogen, then they were stored at −80°C for DNA extraction for sequence analysis. From each generation, five individual plants were sequenced, resulting in a total of fifteen samples for WGS and WGBS analysis.

### Whole Genome Sequencing (WGS) and Whole Genomic Bisulfite Sequencing (WGBS)

The total genomic DNA was extracted from approximately 100 mg of leaf tissue homogenized in liquid nitrogen using a CTAB protocol. The isolated genomic DNA was used for both whole genome sequencing (WGS) and whole genome bisulfite sequencing (WGBS) to assist in identifying the genomic and epigenomic variations. The WGS and WGBS libraries construction and sequencing have been carried out at the Centre d’expertise et de service Génome Québec, Montreal, Canada.

### WGS libraries construction and sequencing

gDNA was quantified using the Quant-iT™ PicoGreen® dsDNA Assay Kit (Life Technologies). Libraries were generated using the NEBNext Ultra II DNA Library Prep Kit for Illumina (New England BioLabs) as per the manufacturer’s recommendations. Adapters and PCR primers were purchased from IDT. Size selection of libraries contained the desired insert size has been performed using SparQ beads (Qiagen). Libraries were quantified using the Kapa Illumina GA with Revised Primers-SYBR Fast Universal kit (Kapa Biosystems). Average size fragment was determined using a LabChip GX (PerkinElmer) instrument.

### WGBS libraries construction

gDNA was quantified using the Quant-iT™ PicoGreen® dsDNA Assay Kit (Life Technologies). Libraries were generated with the NEBNext Ultra II DNA Library Prep Kit for Illumina (New England BioLabs) using 250 ng of input genomic DNA. Adapters were purchased from NEB. Size selection of libraries containing the desired insert size have been performed using sparQ PureMag Beads (Quantabio). Bisulfite conversion had been carried out with the EZ DNA Methylation-Lightning Kit (Zymo Research, Irvine, CA, USA). Libraries were quantified using the Kapa Illumina GA with Revised Primers-SYBR Fast Universal kit (Kapa Biosystems) and average size fragment was determined using a LabChip GX (PerkinElmer) instrument.

### WGS and WGBS sequencing

The libraries were normalized and pooled and then denatured in 0.05N NaOH and neutralized using HT1 buffer. ExAMP was added to the mix following the manufacturer’s instructions. The pool was loaded at 200pM on an Illumina cBot and the flowcell was ran on a HiSeq X for 2×151 cycles (paired-end mode). A phiX library was used as a control and mixed with libraries at 1% level. The Illumina control software was HCS HD 3.4.0.38, the real-time analysis program was RTA v. 2.7.7. Program bcl2fastq2 v2.20 was then used to demultiplex samples and generate fastq reads.

### The computation and analysis of genome sequence data

Raw sequencing reads were quality controlled and trimmed using Trim Galore software (version 0.4.4). The trimmed reads were aligned to the TAIR10 reference genome, and the duplicates were marked using Picard tool. Local realignments around SNPs and INDELs were performed using GATK (genome analysis toolkit) which accounts for genome aligners and mapping errors and identifies the consistent regions that contain SNPs and INDELs. The resulting reads were quality controlled with Haplotype scores and variant sample sites were called individually and jointly using the HaplotypeCaller with GATK. The sites marked as those that had a low-quality score by GATK were filtered out. The effects of variants in the genome sequences were classified using the SnpEff program (Cingolani et al. 2012). According to the SnpEff program (Cingolani et al. 2012) utilized in this study, upstream was defined as 5 kb region upstream of the distal transcription start site, and downstream was defined 5 kb region downstream of the most distal polyA addition site. Variants affecting the non-coding regions were expounded, and biotypes were identified with the available information after comparing with the TAIR 10 reference *Arabidopsis* genome.

The genomation tools were used to obtain a biological understanding of genomic intervals over the pre-defined functional regions such as promoters, exons, and introns and the Functional Classification SuperViewer to create gene association profiles and show the difference between samples. The genes nearest to the non-overlapping SNP and INDEL sites were annotated.

### The computation and analysis of WGBS data

WGBS allows for the investigation of genome-wide patterns of DNA methylation at a single-base resolution. It involves the sodium bisulfite conversion of unmethylated cytosine into uracil, with the resulting cytosine residues in the sequence representing methylated cytosine in the genome which is then mapped to a reference genome (Susan et al. 1994). Binomial tests were applied and used to determine the observed methylation frequency against the bisulfite conversion reaction, and the percentage of methylation levels were calculated at each base (Schultz et al. 2012).

The WGBS raw sequencing data were analysed using tools found in the *methylKit* package. Raw sequencing reads were quality controlled and trimmed using Trim Galore software (version 0.4.4). The trimmed reads were then aligned to the TAIR10 reference genome using the bisulfite mapping tool Bismark (Krueger and Andrews 2011). The methylated cytosines (mCs) were extracted from the aligned reads with the Bismark methylation extractor on default parameters followed by the computation of methylation frequency using the R package software, *methylKit*. The percentage of methylation was calculated by counting the frequency ratio of Cs divided by reads with C or a T at each base and computed at bases with coverage ≥ 10 (Akalin et al. 2012).

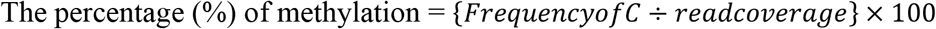

Common bases covered across all samples were extracted and compared, and the differential hyper- and hypo- methylated positions in each chromosome were extracted. The differentially methylated positions (DMPs) overlapping with genomic regions were assessed (in the preference of promotor > exon > intron), and the average percentage of methylation of DMPs around genes with the distances of DMPs to the nearest transcription start sites (TSSs) were also calculated.

The annotation analysis was performed with *genomation* package within *methylKit* to obtain a biological understanding of genomic intervals over the pre-defined functional regions such as promoters, exons, and introns (Akalin et al. 2014). The functional commentary of the generated gene methylation profiles was performed using the SuperViewer tool with Bootstrap to show the difference between samples (Provart and Zhu 2003). Hierarchical clustering of samples was used to analyse similarities and detect sample outliers based on percentage methylation scores. Principal Component Analysis (PCA) was utilized for variations and any biological relevant clustering of samples. Scatterplots and bar plots showing the percentage of hyper-/hypo- methylated bases, the overall chromosome and heatmaps were used to visualize similarities and differences between DNA methylation profiles.

### The Differentially Methylated Regions (DMRs)

A comparison of differential DNA methylation levels between samples reveals the locations of significant differential changes in the epigenome. The obtained information of DMRs was investigated over the predefined regions in all contexts: CG, CHG, and CHH for 100 bp and 1,000 bp tiles across the genome to identify both stochastic and treatment associated DMRs (Akalin et al. 2012).

The differential hyper-/hypo- methylated regions were extracted and compared across the samples. By default, DMRs were extracted with q-values < 0.01 and the percentage of methylation difference > 25%. The differential methylation patterns between the sample groups and the differences in methylation events of per chromosome were also extracted. The methylation profiles of the sample groups used were F25H versus F2C, F25H versus F25C, and F25C versus F2C. In summary, the sliding windows of 100 bp and 1,000 bp were considered for both DMRs and DMPs, and extractions were made based on at least 25% and 50% differences (q-values < 0.01) to assess the significant differences among samples.

### Biological Enrichment Analysis

The functional classification of variants (DMRs and DMPs) unique to each test group was interpreted using SuperViewer to identify regions with the statistically significant number of over- or under-represented genes and genomic features. Biological processes that might be enriched or under-represented within and between generations were assessed. All values were normalized by bootstrap x100, and p-values < 0.05 were retrieved as significant.

### Statistical Analysis and Quality Control Values

The mapped reads were obtained with a quality score of <30, the differential hyper- and hypomethylated bases were extracted with q-values < 0.01 and the percentage of methylation difference larger than 25% in *methylKit.* The Heatmaps of differentially methylated bases were quantified at q-values < 0.01, and the percentage of methylation difference was more significant than 50%. The distances of DMPs to the nearest TSSs obtained from *genomation* at both >25% and >50% of methylation change. The distance between TSSs and DMPs was extracted within +/- 1,000 bp and annotated at DMPs >50% methylation difference. DNA methylation profiles obtained from the *melthylKit* used the obtained pairwise correlation coefficients of methylation levels (in %) and the 1-Pearson’s correlation coefficients for hierarchical clustering of samples. Logistic regression and Fisher’s exact test were used for the determination of differential methylation with calculations of q-values and the Benjamini-Hochberg procedure for the correction of p-values. The T-test for mean difference between groups was calculated with p-values < 0.05. The results of global genome methylation were analysed by one-way ANOVA with the Tukey’s multiple comparisons test using GraphPad Prism version 8.4.2 for Windows. The data are shown as the average percentage (with ±SD) of methylated cytosines from five individual methylomes in each of the progenies. The asterisks show a significant difference between the stressed progeny and the parental control progeny (\**p < 0.1, ** p < 0.05*). The phenotypic data were analysed by the unpaired *t*-test with Welch’s correction using GraphPad Prism version 9.1.1 (255) for Windows. The data were shown as mean ±SD. A *p-value* less than 0.05 (*p ≤ 0.05*) was considered statistically significant.

## Results

### The progeny of stressed plants showed the heat stress tolerance phenotype

The F25H progeny was studied for heat stress memory and response. The response to heat stress was determined at the germination, seedling or mature-plant stage. The F25H stressed progeny did not display any significant difference compared to F25C in germination index represented by the speed of germination and the total germination percentage (Figure 1B). When we exposed 10 days old seedlings of F25C and F25H groups to the same stress that their ancestral progenies were exposed to, the F25H showed slightly higher fresh weight than F25C under heat stress however it was not significantly different (Supp. Figure S1); it should be noted that fresh weight of F25H plants was significantly higher than F25C plants without heat stress. Under heat stress, at the mature-plant stage, the F25H stressed progeny survived better than their respective non-stressed parallel control progenies (p < 0.001) (Figure 1C). The F25H heat-stressed progeny plants also displayed a higher fresh weight when compared with parallel progeny plants.

### The progeny of stressed plants exhibited a higher homologous recombination frequency (HRF)

Our previous reports demonstrated that plants exposed to stress and their progeny exhibit higher HRF (Migicovsky and Kovalchuk 2013; Kathiria et al 2010). Here we also found that the F25H stressed progeny exhibited a higher HRF as compared with parallel control (Figure 1D right panel).

### The multigenerational heat-stressed progeny exhibited higher SNPs and INDELs

Genomic variants consisting of SNPs and INDELs were determined by mapping and comparing F25H, F2C and F25C sequencing reads to TAIR 10 reference genome. The total SNPs identified for F25H, F2C and F25C were 53,678, 15,599 and 15,526, respectively (Figure 2A). The total numbers of INDELs identified for F25H, F2C and F25C were 17,669, 7,986 and 8,143, respectively (Figure 2A). After combining SNPs and INDELs, the total genetic variations in F25H, F2C and F25C were 71,347, 23,585, and 23,669, respectively (Figure 2A). We have also determined the SNPs and INDELs by using stricter parameters, which displayed an overall significantly reduced number of SNPs (12,000 in F25H; 5,190 in F2C; 5,188 in F25C) and INDELs (6,177 in F25H; 4,643 in F2C; 4,781 in F25C), however comparative patterns and conclusion remained almost similar (Supp. Table S1). Relaxed parameters can result in a higher number of false positive genetic variations, and stricter parameters can miss true positives. However, both relaxed and strict parameters outputs resulted in almost similar genetic variations patterns between groups. The parallel F25C and the parental F2C controls exhibited a comparable number of mutations with a slight difference between the numbers of SNPs and INDELs (Figure 2A). Thus, the F25H heat-stressed progeny showed over three times more mutations than the parental and parallel controls (in case of stricter parameters around two times; Supp. Table S1), suggesting that heat-stress induced genetic variations in the stressed progeny (Figure 2A). In addition, INDELs were of a similar size in F25C and F2C, whereas they were significantly larger in F25H (Figure 2B); this is likely caused by multigenerational heat-stress exposure.

**Figure 2.**
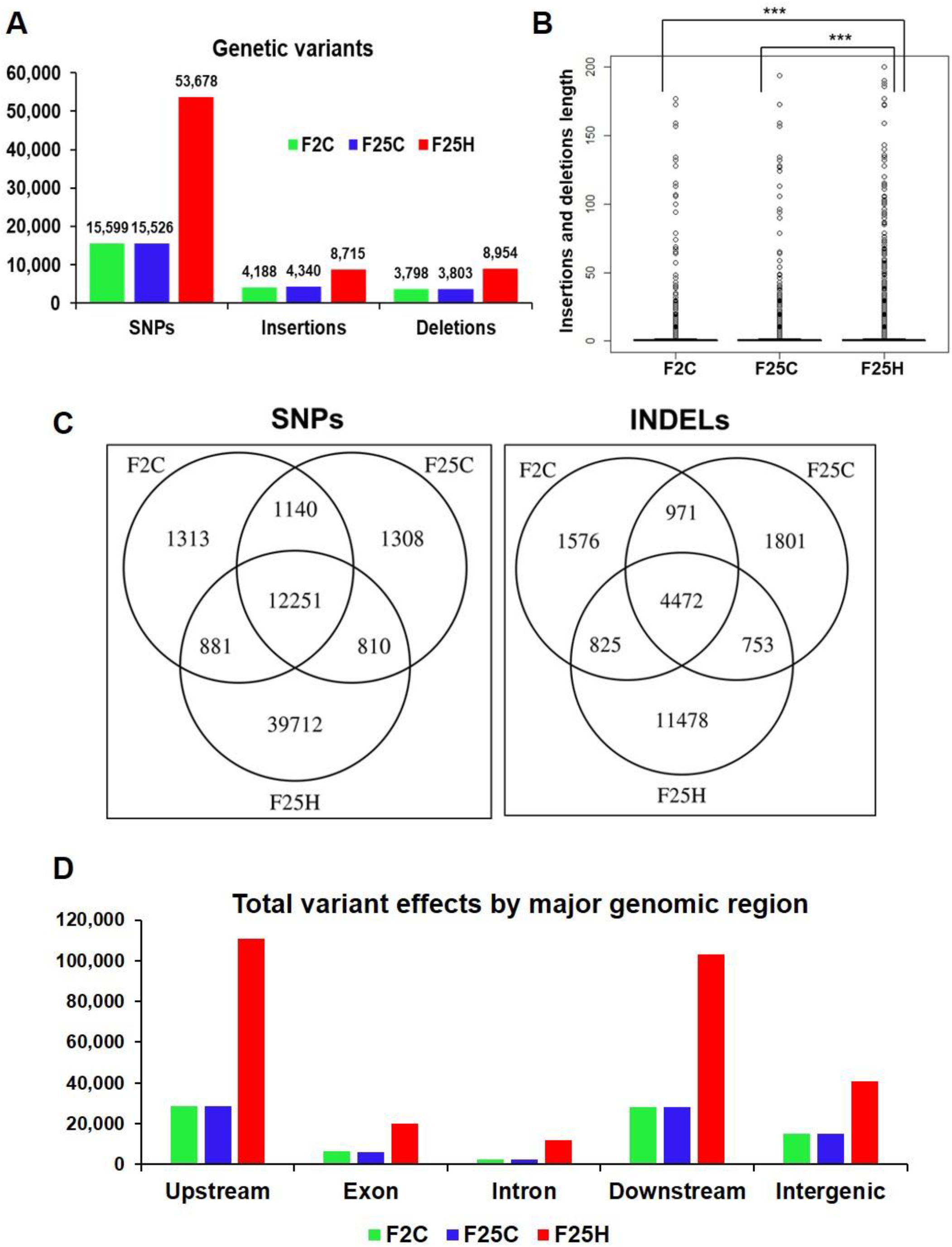
The heat-stressed progeny showed a higher number of genetic variations (with relaxed parameters). (**A**) The total number of SNPs and INDELs in the genomes of stressed progeny and in the parallel and parental control progenies when all samples were jointly considered. Genomic variants consisting of SNPs and INDELs were determined by mapping and comparing F25H, F2C and F25C sequencing reads to TAIR 10 reference genome. (**B**) The box plot for the length of insertions and deletions of variants within F2C, F25C and F25H. The asterisk above bracket (*) shows a significant difference between F25H vs. F2C and F25H vs. F25C where *** indicates *p < 0.001*. Wilcoxon rank-sum text was used to determine a statistical difference (*p<0.05*). (**C**) A Venn diagram of jointly called variants. The non-overlapping area of Venn indicates the number of unique SNPs and INDELs in each sample group. Sample variants were jointly called by the “GATK haplotype Caller” using five samples together. (**D**) The total number of variant effects by major genomic regions. The number of variant effects in the major genomic regions are shown above each bar, and the genomic regions are indicated on the x-axis.

The GATK Haplotype caller calls SNPs and INDELs via an assembly of haplotypes in the active region of the genome which enables the identification of variants that are unique to each sample genome by extracting the non-overlapping sites specific for F25H, F25C, and F2C. The analysis of non-overlapping site-specific variants revealed 39,712, 1,313 and 1,308 unique SNPs for the F25H, F2C, and F25C plants, respectively; 11,478, 1,576 and 1,801 unique INDELs were identified for F25H, F2C and F25C, respectively (Figure 2C). In case of analysis with stricter parameters, we observed an overall significantly reduced number of unique SNPs (7,474 in F25H; 613 in F2C; 629 in F25C) and INDELs (2,467 in F25H; 794 in F2C; 953 in F25C), however comparative patterns remained almost similar to relaxed parameters (Supp. Table S1). Thus, the F25H plants showed a dramatically higher number of unique SNPs and INDELS than the control plants, suggesting that exposure to multigenerational heat predominantly led to the generation of SNPs and INDELs (Figure 2C). The parental F2C and parallel F25C controls showed a very similar number of unique SNPs (Figure 2C).

### The heat-stressed progeny exhibited a higher number of the effects of genetic variants

The total number of genetic variant effects calculated by the corresponding genomic locations of SNPs and INDELs such as the intronic, exonic, untranslated regions (5’ UTR or 3’UTR), upstream and downstream of gene regions and intergenic regions were calculated. Among SNPs, the potential effect on coding sequences was analysed by calculating the number of synonymous or non-synonymous mutations. Positional profiles of SNPs and INDELs revealed a higher number of variant effects in the F25H heat-stressed progeny compared with the non-stressed parallel F25C and parental F2C progenies (Figure 2D). The F25H plants showed the highest total number of variant effects (293,405) compared with the F25C (82,179) and F2C (81,657) controls (Figure 3). The classification of variant effects by genomic regions revealed the largest number of effects observed in upstream of the gene (F25H - 110,837; F25C - 28,762; F2C - 28,403) followed by downstream (F25H - 103,301; F25C - 28,035; F2C - 27,947) and intergenic regions (F25H - 40,941; F25C - 15,184; F2C - 15,058) (Figure 2D, 3). Variant effects were also found within the 5’ untranslated (F25H - 1,817; F25C - 438; F2C - 442) and the 3’ untranslated regions (F25H - 2,312; F25C - 461; F2C - 442) of the genome (Figure 3). It appears that the difference in variants was most drastic in 5’ and 3’ untranslated regions, with ∼4.2- and 5.0-fold difference between F25H and either of the control samples, followed by upstream (∼3.9), downstream (∼3.7) and intergenic regions (∼2.7).

**Figure 3.**
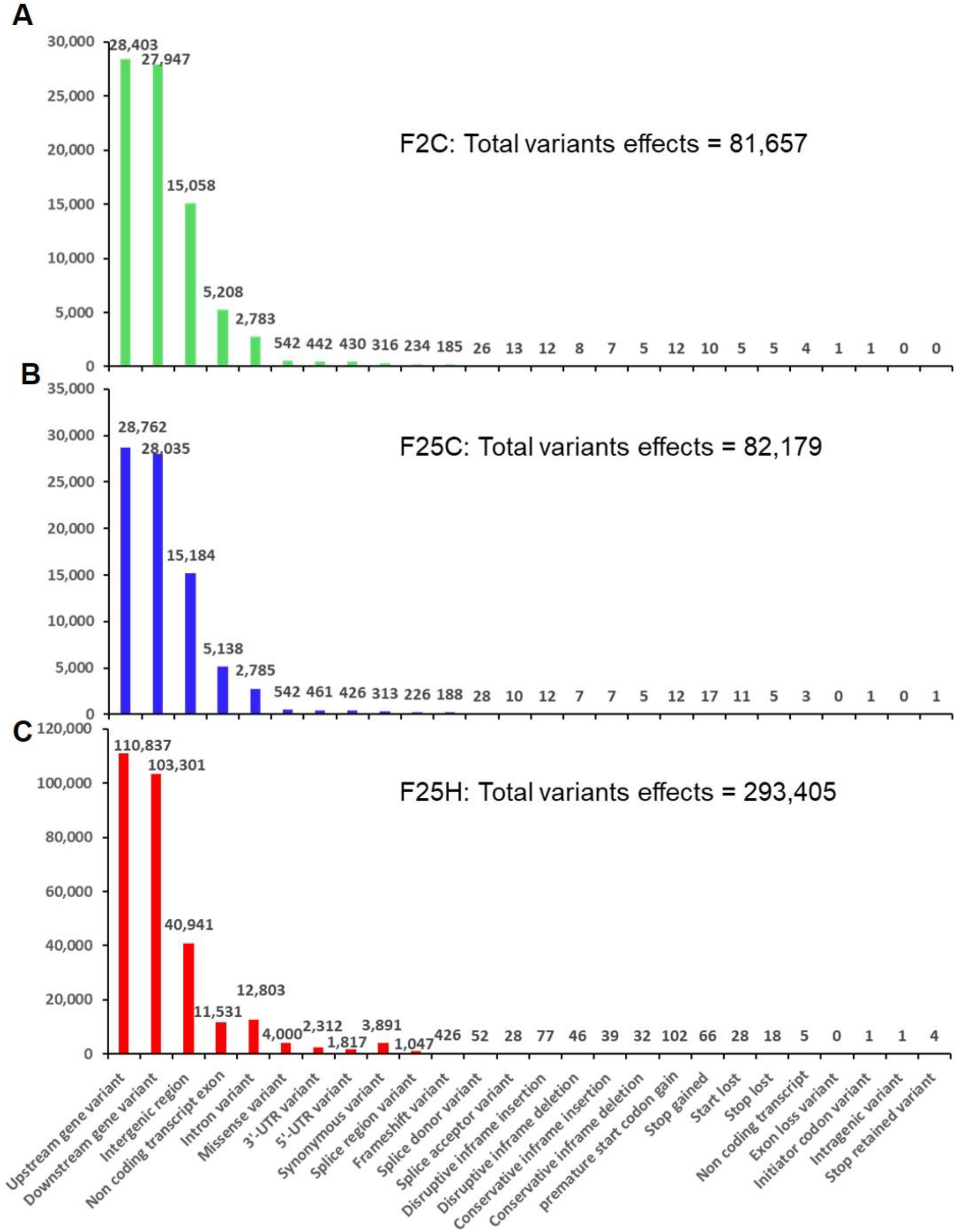
The distribution of variants across the genome by their potential effects based on genomic positions. (A) F25H, (B) F25C and (C) F2C. The number of variant effects is shown above each bar, and genomic positions effects from SnpEff are listed on the x-axis.

The F25H plants showed 12x and 6x higher number of synonymous and nonsynonymous mutations, 3,891 and 4,510, respectively, than the F25C plants (313 - synonymous, 752 – non-synonymous) and the F2C plants (316 - synonymous, 742 – non-synonymous) (Figure 3). In F25H, 18 SNP missense and 66 SNP nonsense mutations were identified (Figure 3C). These numbers were much lower in F25C and F2C where 5 and 5 stop codon losses and 17 and 10 stop codon gains were observed, respectively (Figure 3A, B). The non-synonymous mutations contribute to biological variations in the living organism, consequently, they are subjected to natural selection.

### The analysis of epigenetic variations

#### Multigenerational exposure to heat stress reduces the percentage methylation of total global DNA in the CHG and CHH contexts

The bisulfite sequencing data revealed that the average percentage of global genome methylation in the CG context did not show any significant difference between the F25H plants and either of the control samples (Figure 4A). However, in the CHG context, the F25H plants showed a significantly lower global methylation level (*p < 0.10*) than the control plants (Figure 4A). Similar data were found for the CHH context; the F25H plants exhibited a significantly lower methylation level (*p < 0.05*) compared with the controls (Figure 4A). The reduction of methylation levels in the CHG and CHH contexts might be a part of heat stress adaptation strategy.

**Figure 4.**
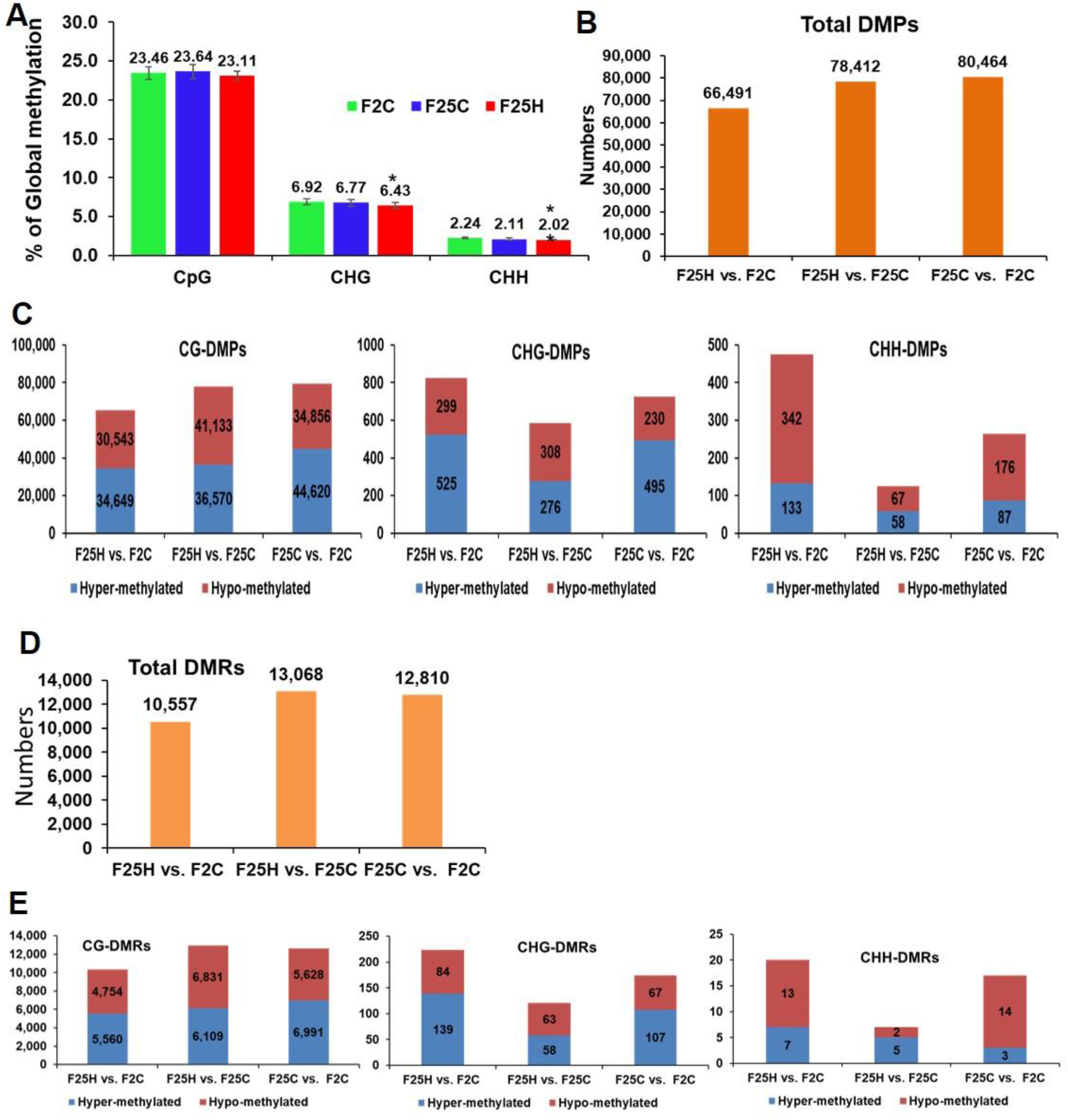
Global methylation changes, differentially methylated positions (DMPs) and differentially methylated regions (DMRs) in F25H, F25C and F2C plants. (**A**) The average percentage of methylated cytosines in F2C, F25C, and F25H in the CG, CHG, and CHH sequence contexts (H=A, T, C and mean, n=5). Methylation levels were determined from reads with the minimum coverage ≥ 10 mapped to the TAIR 10 reference genome by using Bismark software. The data were analysed by one-way ANOVA with the Tukey’s multiple comparisons test using GraphPad Prism version 8.4.2 for Windows. The data are shown as the average percentage (with ±SD) of methylated cytosines from five individual methylomes in each of the progenies. The asterisk shows a significant difference between the stressed progeny and the parental control progeny (\**p < 0.1, ** p < 0.05*). (**B**) The total number of DMPs in F25H vs. F2C, F25H vs. F25C and F25C vs F2C. (**C**) The total number of DMPs in the CG, CHG and CHH contexts in F25H vs. F2C, F25H vs. F25C and F25C vs F2C. (**D**) The total number of DMRs in F25H vs. F2C, F25H vs. F25C and F25C vs F2C comparison groups. (**E**) The total number of DMRs in CG, CHG and CHH contexts in F25H vs. F2C, F25H vs. F25C and F25C vs F2C.

#### The analysis of the number of DMPs and DMRs reveals a higher number of differentially methylated cytosines in the CHG and CHH contexts in the F25H plants

The total number of DMPs for F25H vs. F2C, F25H vs. F25C and F25C vs. F2C were 66,491, 78,412 and 80,464, respectively, suggesting that there is no effect of heat stress on the total number of DMPs (Figure 4B). The analysis of DMPs in a specific sequence context revealed a different pattern. While DMPs in the CG context were higher in the F25C vs. F2C control comparison group, in the CHG and CHH contexts, DMPs were much higher in comparison groups involving the F25H stressed progeny (Figure 4C). The difference was especially obvious for hypomethylated cytosines at the CHG context; the F25H vs. F2C group showed 299 hypomethylated DMPs, while the F25C vs. F2C group - only 230. A similar pattern was observed in the CHH context, nearly a 2-fold larger number of hypomethylated DMPs was observed in the F25H vs. F2C group in comparison with the F25C vs. F2C group - 342 vs. 176, respectively; the difference for hypermethylated DMPs was also substantial, 133 vs. 87 (Figure 4C). These data suggest that epimutations induced by multigenerational heat stress are primarily associated with changes in the CHG and CHH contexts, and that hypomethylation is likely a prevalent mechanism.

The total DMRs in the100 bp window for F25H vs. F2C, F25H vs. F25C and F25C vs. F2C were 10,314, 12,940, 12,619, respectively (Figure 4D). In the case of the 1,000-bp window, the total DMRs for F25H vs. F2C, F25H vs. F25C and F25C vs. F2C were 49, 36, 32, respectively. As to the CHG and CHH context, there was a larger number of DMRs in the F25H vs. F2C group compared with other groups (Figure 4E).

### The progeny exposed to multigenerational heat-stress did not show any difference in the methylation levels

The percentage of methylation (the analysis of the percentage of methylated cytosines at specific sites) in F2C, F25C, and F25H showed that most of the cytosines had either a high (70-100%) or a low (0-10%) level of methylation in the methylated CG context, however we did not see any difference in methylation levels between F2C, F25C, and F25H progeny (Supp. Figure S2). Cytosine methylation in the CHG context was much lower compared with the CG context; most sites had 0-60% methylation levels including very high number of cytosines methylated at 0-10%, and sites with high levels of methylation (70-100%) were not observed (Supp. Figure S3). Similarly, methylation in the CHH context was much lower than in the CG or CHG contexts; there was an even lower frequency of occurrence of sites with the 0-20% methylation level (Supp. Figure S4). When each sequence context was considered, there was no significant difference in the methylation levels between the F2C, F25C, and F25H groups. In general, F2C, F25C, and F25H showed a similar percentage of methylation in the CG, CHG, and CHH contexts (Supp. Figure S2-4).

### The heatmaps based hierarchical clustering analysis of DMPs and DMRs revealed directional changes of heat-induced epimutations in the stressed progeny

We analyzed the relatedness of the control and stressed progeny samples using the hierarchical clustering heatmap analysis (Figure 5). The heatmap analysis of changes in both hypo- and hyper-methylated DMPs revealed a clear separation of five samples of F25H from five samples of the F2C parental progeny in all three CG, CHG and CHH cytosine methylation contexts (Figure 5A). Similarly, a comparison between F25H and F25C also showed separate clustering (Supp. Figure S5A). A comparison between F25C and F2C samples showed more similarity, with two F25C samples clustering together with F2C samples (Supp. Figure S5B). Heatmap clustering of CG-DMRs also revealed a clear separation between F25H and F2C samples (Figure 5B). Considering that CHH methylation happens in plants *de novo*, separate clustering of stressed samples from control progeny suggests the importance of CHH methylation in stress adaptation and indicates that multigenerational exposure to heat stress likely induces non-random changes in the methylome in the CHH context.

**Figure 5.**
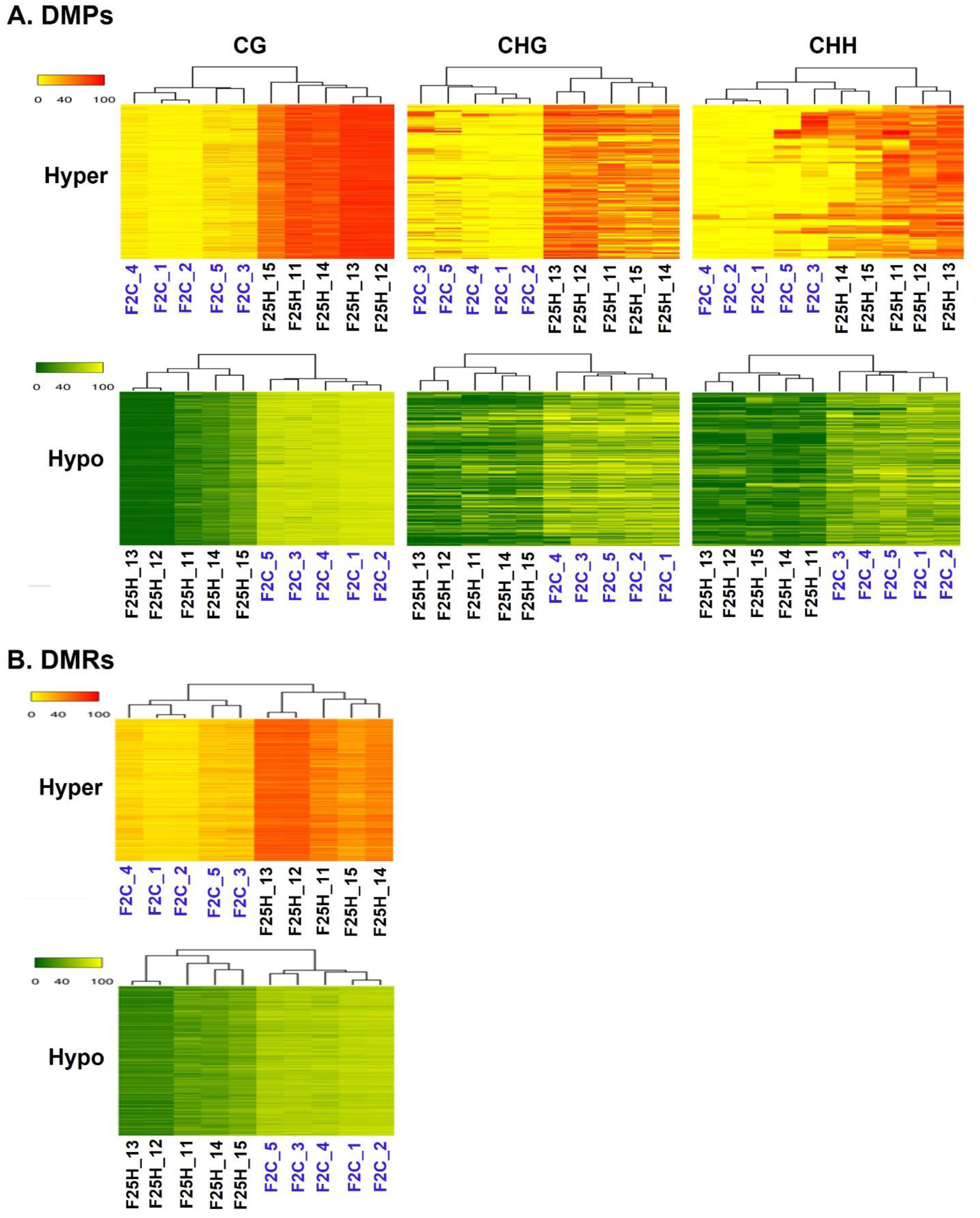
A hierarchical clustering heatmap analysis. (**A**) Heat maps of DMPs for hypermethylated cytosines (the upper panel) and hypomethylated cytosines (the lower panel) in CG, CHG and CHH contexts in F25H vs. F2C. Differentially methylated cytosines in the genome with differences of > 50% in F25H vs. F2C. (**B**) Heat maps of DMRs of hypermethylated cytosines (the upper panel) and hypomethylated cytosines (the lower panel) in CG contexts in F25H vs. F2C. Differentially methylated positions in the genome with differences > 50% \ in F25H vs. F2C. In ‘the upper panel’, the red section indicates the larger percentage of methylation, and the yellow section indicates the lower percentage, and in ‘the lower panel’, the green section indicates the larger percentage of methylation and the yellow one indicates the lower percentage, q-value <0.01.

### Annotation of genes with DMPs and DMRs

DMPs and DMRs were then characterized to determine whether they were preferably located near genes. The location of hypo- or hyper-methylated DMRs was compared to the annotated Arabidopsis genes using *genomation* Bioconductor package (Akalin et al., 2014). Both hyper- and hypo-methylated CG-DMRs were predominantly located in the gene body, while were less frequent at TSS (Transcription Start Site) and TTS (Transcription Termination Site) regions (Figure 6A, C). In contrast, in the case of TEs, F25H progeny displayed less pronounced hypermethylation in CG-DMRs located in TEs body as compared to upstream and downstream regions of TEs (Figure 6B, D). Overall, methylation changes in the gene body were much more pronounced than in TEs in F25H group as compared to other groups (Figure 6).

**Figure 6.**
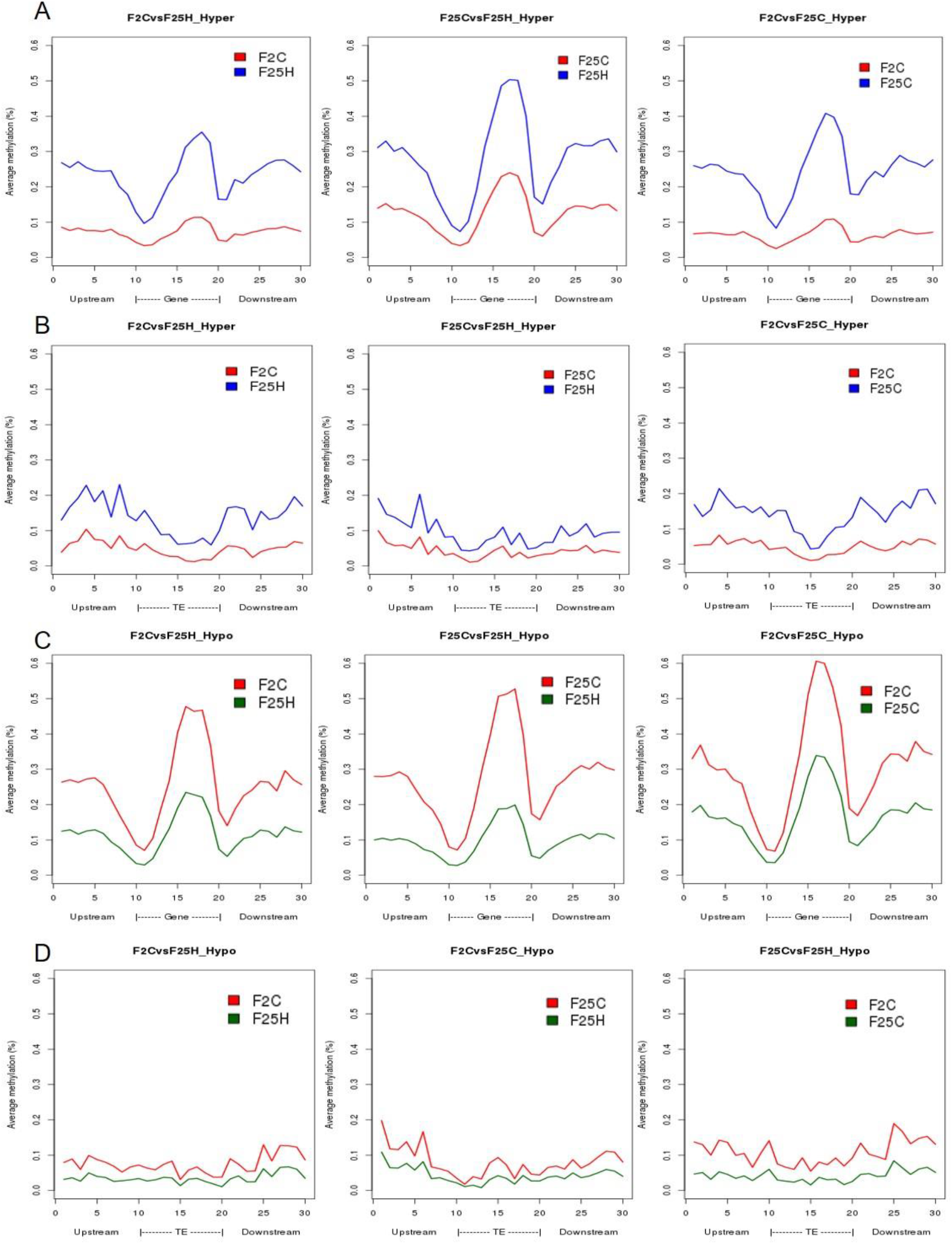
>The distribution of hyper-and hypo-methylated CG-DMRs in the gene body and TEs in F25H vs. F2C, F25H vs F25C and F25C vs. F2C comparison groups. (**A**) Distribution of hypermethylated CG-DMRs in gene bodies. (**B**) Distribution of hypermethylated CG-DMRs in TEs. (**C**) Distribution of hypomethylated CG-DMRs in gene bodies. (**D**) Distribution of hypomethylated CG-DMRs in TEs.

In the CG context, the percentage of hypermethylated and hypomethylated DMPs in all groups (F25H vs. F2C, F25H vs. F25C and F25C vs F2C) was the highest in the exon region, followed by the promoter, intron and intergenic regions, (Figure 7A-F). In the case of the CHG context, all comparison groups (F25H vs. F2C, F25H vs. F25C and F25C vs F2C) showed the highest percentage of hypermethylated DMPs in the promoter, followed by intergenic, exon and intron regions (Figure 7A, C, E). Interestingly, F25H vs. F2C groups showed the highest percentage of hypomethylated DMPs in the intergenic regions (48%) followed by the promoter (44%), while in the F25C vs F2C group, the highest percentage of DMPs was in the promoter region (52%) followed by the intergenic regions (44%) (Figure 7F). In the CHH context, in all groups, the percentage of both hyper- and hypo- DMPs was higher in the promoter, followed by the intergenic, exon and intron regions (Figure 7).

**Figure 7.**
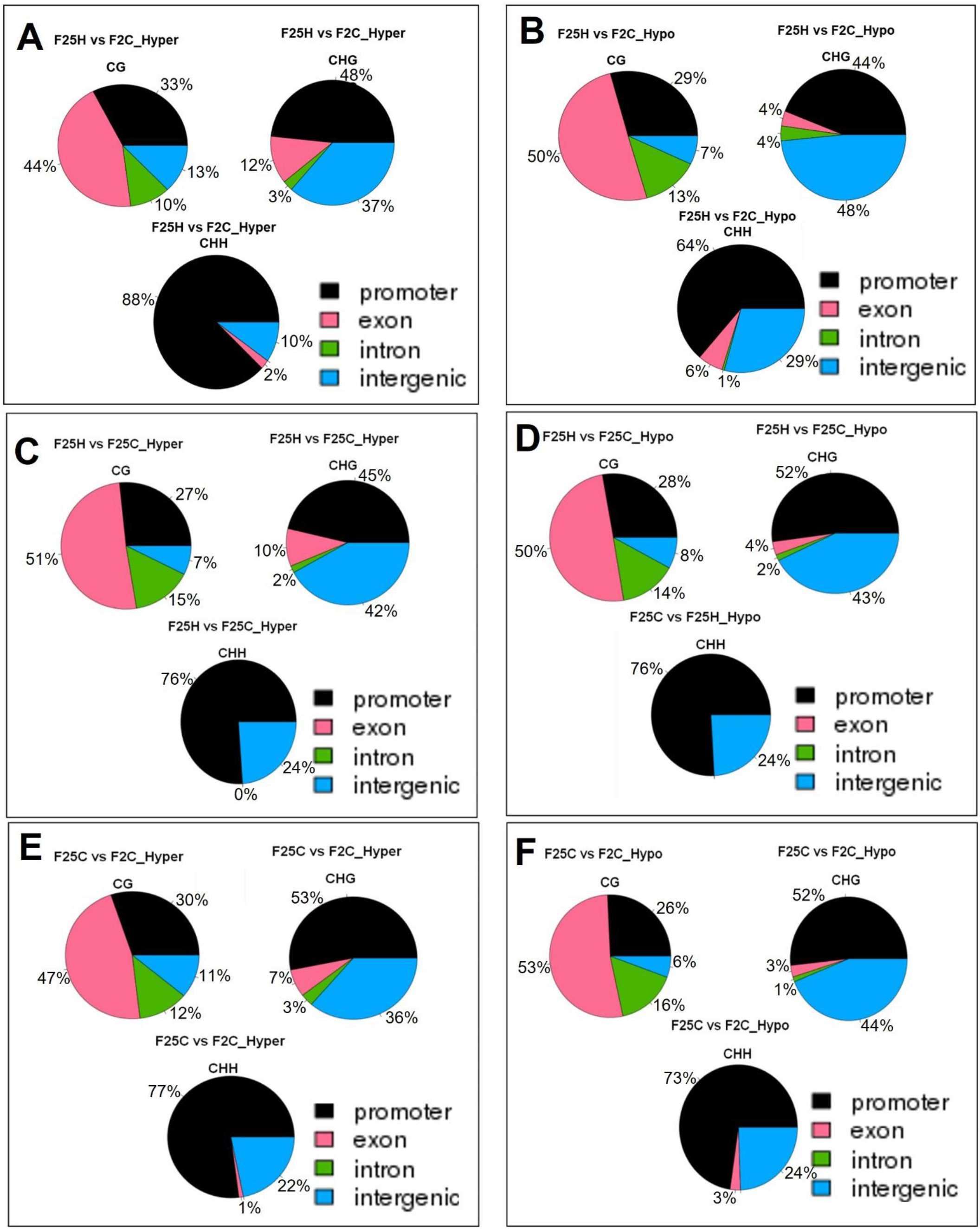
The distribution of hyper-and hypo-methylated DMPs in the genome in the comparison groups F25H vs. F2C (A – hyper, B – hypo), F25H vs F25C (C – hyper, D – hypo) and F25C vs. F2C (E – hyper, F – hypo). The data are shown as an average percentage of differentially methylated cytosines at the promoter, exon, intron and intergenic regions. DMPs were considered as regions of methylation with > 25% differences.

Overall, in the case of the CG context, the highest percentage of DMPs was observed in the gene body, whereas in the case of the CHG and CHH contexts, it was highest in the promoter and intergenic regions.

### Comparative analysis revealed the heat-inducible nature of genetic variants and the spontaneous nature of epigenetic variants

Strict genetic analysis showed that the ratio between the number of epigenetic and genetic variants was 6.27x, 7.3x and 35.9x in the comparison groups of F25H vs F2C, F25H vs F25C and F25C vs F2C, respectively (Table 1), indicating that epigenetic variations were more prevalent than genetic variations. The largest difference in the ratio of epigenetic to genetic variations was observed between F25C and F2C control plants, which indicated the spontaneous nature of epigenetic variations. In the case of epigenetic variations, F25C parallel control progeny showed 1.2 times (80,464 vs 66,491) more variation than F25H stressed progeny when both were compared with the parental control progeny F2C. In contrast, the F25H stressed progeny displayed 5 times (10,597 vs 2,238) more genetic variations than the F25C parallel control progeny when both were compared with the parental control progeny F2C. These comparisons suggest that heat stress predominantly leads to the accumulation of genetic rather than epigenetic variations.

**Table 1.** Comparison of epigenetic and genetic variations in F25H, F25C and F2C plants.

| Samples group | Epigenetic variants | Genetic variants by using relaxed parameters | Genetic variants by using strict parameters | Epigenetic vs. Genetic variants (with strict parameters) |
| --- | --- | --- | --- | --- |
|  | Total DMPs | Total (SNPs+INDELs) | Total (SNPs+INDELs) |  |
| **F25H vs F2C** | 66,491 | 52,753 | 10,597 | 6.27x |
| **F25H vs F25C** | 78,412 | 52,896 | 10,636 | 7.3x |
| **F25C vs F2C** | 80,464 | 4,672 | 2,238 | 35.9x |

### There was no difference in the enrichment of genetic variants for any biological process in F25H

The most enriched biological process for SNPs was classed as ‘unknown biological processes’ regardless of the tested group (Figure S6). The only over-represented group was ‘transport’ found in F2C INDELs sample. All other biological processes were underrepresented for all samples, and no significant difference between the F25H group and any other group was observed (Supp. Figure S7, Table S2).

### Epimutation-associated genes participate in stress response pathways

Gene Ontology (GO) analysis of epimutations (DMRs) in the hypermethylated CG context in F25H group showed a significant difference in the enrichment of biological processes such as ‘response to abiotic or biotic stimuli’, ‘cell organization and biogenesis’ and ‘DNA or RNA metabolism’ compared to F2C and F25C groups (Figure 8; Table S3). In the hypomethylated CG context, no significant difference was observed between the F25H vs F2C group and other groups (Supp. Figure S9; Table S4).

**Figure 8.**
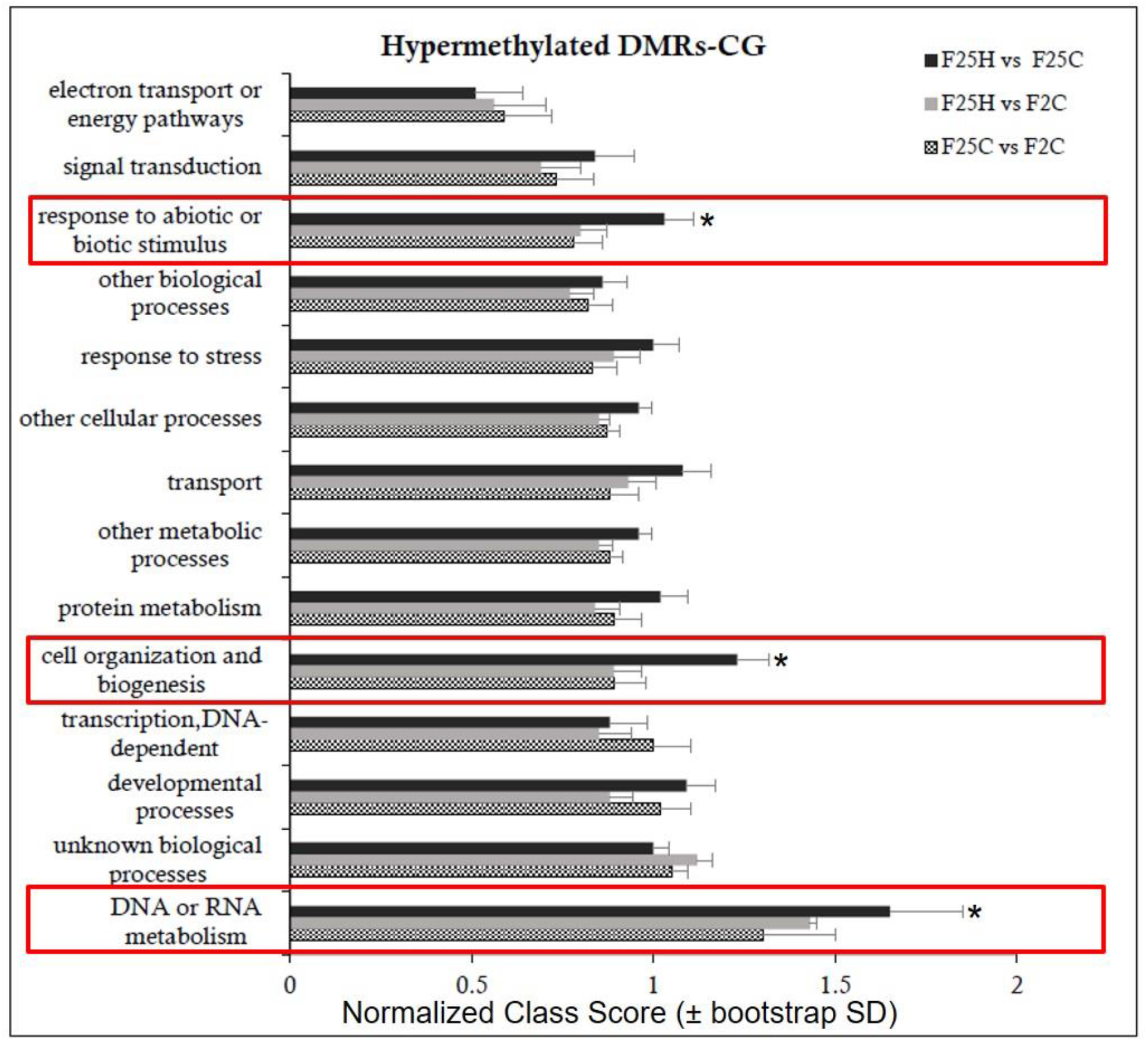
Enrichment analysis of hypermethylated DMRs on CG sites and their classification based on biological processes. Y-axis shows normalized Class Score with binomial coefficients as calculated by SuperViewer. To calculate enrichment, * p-value of < 0.05, ±bootstrap StdDev were used.

In the hypomethylated CHG context, the F25H vs F2C group exhibited the enrichment in ‘protein metabolism process’ which was not observed in other groups (Supp. Figure S9A; Table S5). In the hypomethylated CHH context, the F25H vs F2C group exhibited the enrichment in ‘transport’, ‘other metabolic processes’, and ‘developmental’ processes (Supp. Figure S9B; Table S6).

## Discussion

The current study aimed to understand the effects of multigenerational exposure to heat stress on plant phenotypes, genotypes, and epigenotypes. Specifically, we compared the progeny of plants exposed to heat stress over 25 consecutive generations with their counterparts (the parental and parallel control progenies). We studied the genetic variations (SNPs and INDELs) by using the whole genome sequencing analysis as well as the epigenetic variations (DMPs and DMRs) by using the whole genome bisulfite sequencing analysis of the heat-stressed and control progenies. Overall, this study has deciphered the phenotypic, genomic and epigenomic changes in the heat-stressed progeny in response to multigenerational exposure to heat stress.

### Multigenerational heat stress induced the phenotypic resilience in the stressed progeny

The phenotype of the progeny can be affected by the parental environment (Groot et al. 2016). Multigenerational stress exposure elicits different outcomes compared to single generation exposure (Groot et al. 2016). Our study showed that the progeny derived from 25 generations of heat stress displayed a notable stress tolerance at the mature stage (Figure 1C). Like our study, Wibowo et al. (2018) reported that the direct progeny of G2-G5 stressed plants exhibited the higher germination and survival rates under high-salinity stress. The authors concluded that stress priming requires repetitive exposure to stress. In some cases, it was evident that parental heat stress affected the response of offspring generation because heat stress accelerated leaf production to speed up the development of the progeny of stressed plants (Liu et al. 2015; Porter 2005). Our previous studies showed that the heat-induced modifications of gene expression might have a role in phenotype changes such as leaf number and flowering time (Migicovsky et al. 2014). Zheng et al. (2017) also found that multigenerational exposure to drought improved the drought adaptability of rice offspring. The analysis of plant weight is useful for determining the active growth and development of a plant (Yadav et al. 2012). F25H heat-stressed progeny plants displayed a higher fresh weight when compared with parallel progeny plants.

The degree or severity of high temperature stress can impact cellular homeostasis (Kotak et al. 2007), and exposure to above the optimum growth temperature over time can result in the acquired thermotolerance (Bray 2000; Sung et al. 2003). Moreover, the expression of transgenerational effects in the form of changes of phenotype responses strongly depends on the offspring environment (Boyko and Kovalchuk 2010; Groot et al. 2016) and varies between the genotype and trait (Suter and Widmer 2013a; Verhoeven and van Gurp 2012). Some reports have mentioned the effects of epigenetic inheritance or transference of transgenerational stress memory to be restricted to a single generation without the maintenance of stress (Pecinka et al. 2009; Suter and Widmer 2013b). Also, it was shown that multigenerational exposure often reduces the expression of parental effects compared to single-generation exposure (Groot et al. 2016).

Both intergenerational and transgenerational changes in response to stress include alterations in homologous recombination frequency (HRF). The progeny of plants that were stressed show the elevated HRF (Migicovsky and Kovalchuk 2013). In the current report, under non-stress conditions, the stressed progeny F25H exhibited the elevated HRF compared with their parallel control progenies (Figure 1D). It has been reported that plants exposed to salt and UV stresses show a much higher HRF than control plants (Boyko et al. 2010). The progeny of pathogen-infected plant has a higher level of somatic HRF, and the progeny also show the decreased methylation at loci for resistance to tobacco mosaic virus (TMV) and they have global hypermethylation (Boyko et al. 2007). Likewise, the progeny of plants exposed to heavy metal stress showed high HRF (Rahavi et al. 2011). If stresses persist through multiple generations, it is possible that the epigenetic memory will be stronger and persist longer in the absence of stress, thus exhibiting changes in HRF for a longer period of time (Migicovsky and Kovalchuk 2013).

### The analysis of genomic variations induced by multigenerational exposure to heat stress

In our study, we have shown that the progeny of heat-stressed plants (F25H) had 3x higher frequency of genomic variations than the non-stressed control plants. It is well established that environmental stresses can be mutagenic and can cause genome instability (Boyko et al. 2010; Boyko and Kovalchuk 2011; Gill et al. 2015). Previously, Jiang et al. (2014) reported around 2x more genetic variations in 10^th^ generation of salinity stress lineages compared to control lineage. Environmental changes may also increase homologous recombination events or can facilitate the mobilization of transposable elements (Long et al. 2009; Boyko et al. 2010; Migicovsky and Kovalchuk 2013). Eventually, environmental conditions may induce mutations and increase the chances of genomic diversity leading to the adaptation to cope up with the ever-changing environmental conditions (Filichkin et al. 2010).

The position of SNPs may influence the rate of transcription directly or indirectly depending on their corresponding genomic location; SNPs can also directly alter the coding sequence leading to a synonymous or non-synonymous amino acid replacement (Cingolani et al. 2012). SNPs and INDELs are vital genetic variations that can directly disrupt gene function and affect plant adaptability in the changing environment (Shastry 2009). For instance, SNPs can affect light response and flowering time by changing amino acids in phytochromes A and B (Filiault et al. 2008; Maloof et al. 2001). Jain et al. (2014) indicated that INDELs occur in different genomic regions, and they further suggested that positions of INDELs affect the function and expression of genes in a different manner. In the current study, genetic variants were mostly found in the upstream and downstream region of the gene and were significantly higher (3x) in the F25H stressed progeny than in F25C parallel and F2C parental control progenies (Figure 2A, D). In contrast, the number of genetic variations was almost similar between the F25C parallel and F2C parental control progenies (Figure 2A). Very recently, Belfield et al. (2020) also showed higher mutation frequency in 11^th^ generation of mild heat stress lineage (29°C) than control lineage. Positional profiles of genetic variations in our work revealed the higher number of variant effects in the F25H heat-stressed progeny compared with the non-stressed parallel F25C and parental F2C progenies (Figure 3). The F25H heat-stressed progeny exhibited higher rate of synonymous (12x) and nonsynonymous (6x) mutations than the parallel and parental control progenies (Figure 3). SNPs can create new splice sites and alter gene functions (Guyon-Debast et al. 2010). Therefore, the higher number of genetic variations in the heat-stressed progeny may be an indication of the adaptive processes occurring in the progeny of stressed plants.

Interestingly, the INDELS were of a similar size in F25C and F2C, whereas they were significantly larger in F25H (Figure 2B), suggesting that multigenerational exposure to heat stress resulted in an increase in the size of INDELs in the heat-stressed progeny. Larger deletions may have a greater effect on gene expression and changes in the phenotype. It has been reported that small and large INDELs can cause pathogen sensitivity (Mindrinos et al. 1994; Kroymann et al. 2003).

### The analysis of epigenetic variations induced by multigenerational exposure to heat stress

The mechanism behind intergenerational and transgenerational stress memory in plants likely includes changes in the DNA sequence, DNA methylation and chromatin structure (Crisp et al. 2016; Zhang et al. 2018; Perez and Lehner 2019; Lind and Spagopoulou 2018; Miryeganeh and Saze 2020). The inheritance of changes in DNA methylation is one of the mechanisms of adaptation – epialleles often lead to the appearance of new alleles (Becker et al. 2011; Paszkowski and Grossniklaus 2011; Zhang et al. 2018). It has been reported that abiotic stresses can cause hyper- or hypomethylation in the genome after either short-term or long-term exposure (Uthup et al. 2011; Zhang et al. 2018). Several studies suggested that multigenerational stress exposure could cause substantially higher heritable epigenetic variations compared with single-generation exposure (Remy 2010, Groot et al. 2016, Zheng et al. 2017). This phenomenon was explained by the gradual acclimatization of epigenetic effects (Groot et al. 2016).

In our study, the bisulfite sequencing data revealed that the average percentage of global genome methylation in the CG context was similar among all three groups (Figure 4A). However, in the CHH contexts, the F25H stressed progeny showed a lower global methylation than the control progenies (Figure 4A). The data suggest that the reduction of methylation in the CHG and CHH contexts might be a part of adaptation strategies to heat stress. In plants, *de novo* DNA methylation at all three cytosine sequence contexts is established by RNA-directed DNA methylation (RdDM) pathway via DRM2 (DOMAINS REARRANGED METHYLTRANSFERASE 2) methyltransferase that is directed to the target loci by siRNAs (mainly small 24 nucleotide short interfering RNAs). Once the DNA methylation is established, it is maintained by various methyltransferases, including MET1 (METHYLTRANSFERASE 1; mainly CG context), CMT3 (CHROMOMETHYLASE 3; mainly CHG context), and DRM2 (mainly CHH context) (Law and Jacobsen 2010; Stroud et al 2013). Thus, CHH methylation is exclusively established and re-established by the *de novo* RdDM pathway (Erdmann and Picard 2020; Wassenegger and Dalakouras 2021). Therefore, upon meiosis/mitosis, and in the absence of RNA triggers, CG and CHG methylation can be maintained by MET1 and CMT3, respectively, while CHH methylation is lost. In the current study, the global CG methylation level was almost unaffected in F25H stressed progeny, but the global CHH methylation level was significantly reduced, which strongly suggests that under heat stress, the RdDM pathway was affected however, the MET1 pathway was not impaired. It has been reported that abiotic stresses can cause hyper or hypomethylation in the genome after either short-term or long-term exposure (Thiebaut et al. 2019; Uthup et al. 2011). High temperature (HT) induced a global disruption of DNA methylation in cotton anther, mainly causing CHH hypomethylation in HT-sensitive cotton varieties (Ma et al. 2018). A decrease in global DNA methylation might result in the interruption of glucose- and reactive oxygen species (ROS)-producing metabolic pathways which may lead to microspore sterility (Ma et al. 2018). Min et al. (2014) also reported that in cotton anthers, HT decreased the expression of S-ADENOSYL-L-HOMOCYSTEINE HYDROLASE1 (SAHH1) and methyltransferases DRM1/DRM3, resulting in the genome-wide hypomethylation. Another report revealed that in soybean, heat stress induced the global DNA hypomethylation, especially in the CHH context, in both root hairs and stripped roots (Hossain et al. 2017). In cultured microspores of *Brassica napus* cv. Topas, heat-shock treatment (32°C for 6h) triggers DNA hypomethylation, particularly in the CG and CHG contexts (Li et al. 2016). However, Gao et al. (2014) reported that under heat stress (37°C for 2h, and then 45°C for 3h), global genome methylation levels were significantly higher in the heat-sensitive genotype of rapeseed than in the heat-tolerant genotype. In *Brassica rapa,* the global DNA methylation levels are relatively stable under heat stress, while changes in CHH methylation at TE suggest that CHH methylation may be important for heat-stress response and tolerance (Liu et al. 2018). The methylome analysis revealed that water deficit was associated with a decrease in CHH methylation in apple cultivars, which might result in the hypomethylated status of TEs (Xu et al. 2018). Wibowo et al. (2016) also showed that methylation changes in CHG and CHH were well correlated with hyperosmotic stress treatment, whereas changes in CG methylation did not show any correlation, suggesting that CG methylation patterns occurred stochastically in the stressed and non-stressed samples. Further, they confirmed that hyperosmotic stress directed DNA methylation changes primarily at non-CG sites, and these epigenetic modifications were associated with an acquired transient adaptation to stress (Wibowo et al. 2016). Exposure to abiotic stressors such as salt also causes DNA hypomethylation in the progeny; Jeremias et al. (2018) reported the inheritance of DNA hypomethylation in response to salinity stress in *Daphnia magna*. Also, Jiang et al. (2014) showed that lineages exposed to soil salinity stress accumulated more methylation at the CG sites than the control progenies. Recently, Atighi et al. (2020) showed a massive and genome-wide hypomethylation as a crucial plant defence mechanism in response to nematode or bacterial pathogen infection in rice and tomato. Further, they demonstrated that DNA hypomethylation in the CHH context was associated with a reduced susceptibility to root-parasitic nematodes in rice.

Hierarchical clustering of epimutations and the heat map analysis of changes in both hypo- and hypermethylated DMPs revealed that the F25H progeny of stressed plants showed a clear separation from plants of the parental and parallel control progenies (Figures 5, S4), indicating directional epimutations in the F25H stressed progeny due to multigenerational exposure to heat stress. Ganguly et al. (2017) also found similar clustering patterns in response to heat. They showed that in Arabidopsis, the progeny of stressed plants clustered separately from the progeny of non-stressed plants in response to the treatment with heat stress (Ganguly et al. 2017). Becker et al. (2011) compared global DNA methylation patterns among 10 *Arabidopsis thaliana* lines propagated for 30 generations from a common ancestor. Hierarchical clustering based on DMPs grouped siblings of the 3rd and 31st generation lines together in separate groups and suggested that epimutations did not distribute randomly throughout the genome (Becker et al. 2011). Similar to our study, Zheng et al. (2017) showed that the multigenerational drought stress induced directional epimutations in the methylome of rice offspring. Eichten and Springer (2015) reported that the separate hierarchical clustering of epimutations was associated with cold stress treatment.

We observed that the total number of differentially methylated positions (DMPs) was higher between the control groups F25C vs. F2C (80,464), compared with the stressed progeny groups F25H vs. F2C (66,491) and F25H vs. F25C (78,412) (Figure 4B, C). It was somewhat surprising because we expected an opposite result; we hypothesized that heat stress would induce epimutations and expected to see a higher number of epigenomic variations in the progeny of heat stressed plants compared with the controls. It is possible that heat stress does not result in an overall increase in epimutations, but it rather causes epimutations to occur at the specific, heat stress-related sites in the genome. The fact that the observed progenies of heat-stressed plants cluster together confirms this hypothesis. Becker et al. (2011) have compared the genome wide DNA methylation patterns between the 31^st^ generation (that grew under control conditions) and the ancestor generation (the 3^rd^) and deciphered 30,000 differentially methylated cytosines (DMPs). More than one third of recurrent events were observed in the whole DMPs and DMRs, indicating the non-random nature of these epimutations. Further, they reported an average of 3,300 DMPs between immediate generations; this was around three times more than what would have been expected from 30,000 DMPs accumulated between the 3^rd^ and the 31^st^ generation lines (Becker et al. 2011). This data suggests that the number of epimutations does not increase linearly with time, indicating that only a fraction of new DMPs is maintained by the transgenerational inheritance over the longer term (Becker et al. 2011).

The analysis of DMPs in a specific sequence context revealed a different pattern. While DMPs in the CG context were higher in the F25C vs. F2C control group, in the CHG and CHH context, DMPs were higher in comparison groups involving the F25H stressed progeny (Figure 4C). The F25H vs. F2C comparison group showed the higher number of hypomethylated and hypermethylated DMPs in the CHG and CHH contexts compared with the control comparison groups (Figure 4C). These data suggest that the epimutations induced by multigenerational exposure to heat stress are primarily associated with changes in the CHG and CHH contexts, and that hypomethylation is a prevalent mechanism. A similar phenomenon has been narrated in response to heavy metal stresses where it has been reported that hypomethylation prevails at several loci in hemp and clover (Aina et al. 2004). In epigenetic mutants, methylation patterns of DMPs link DNA methylation to the response to the environmental signals like heat stress (Popova et al. 2013) and salt stress (Yao et al. 2012). Several reports also suggest that it is common for plants to methylate cytosines in the sequence contexts of CHG and CHH, but the reestablishment of epigenetic modifications is mostly guided by small RNAs or the heterochromatin-directed methylation pathways (Feng, Jacobsen et al. 2010, Stroud et al. 2013). Further, we observed that in the case of the CG context, the highest percentage of DMPs was observed in gene body, whereas in the case of the CHG and CHH contexts, it was the highest in the promoter and intergenic regions. Changes in the CG sequences were usually found to cluster around the regulatory region of genes (Zheng et al. 2013), while changes in CHG and CHH in the gene body, likely contributing to changes in phenotypes (Bewick and Schmitz 2017). It was reported before that the promoters of stress-responsive genes may be hypomethylated under stress conditions (Yao and Kovalchuk 2011, Bilichak et al. 2012).

In our study, F25H stressed progeny displayed more pronounced methylation changes in gene body while less pronounced changes in the body of TEs (Figure 6). This is an interesting finding, as it is believed that TEs are quite unstable and are activated in response to stress. It remains to be shown whether differences in methylation of TEs in these groups would also result in lower transposon activity in F25H plants. Our previous work demonstrated that the immediate progeny of heat-stressed plants showed higher activity of TEs (Migicovsky et al. 2014). It is possible that multigenerational exposure to heat stabilizes transposons. Indeed, previously, Becker et al. (2011) also reported the transgenerational maintenance of CG methylation of TEs in response to propagation of Arabidopsis for 30 generations under normal conditions.

The Gene Ontology analysis identified that epimutations in the F25H stressed progeny showed mainly enrichment in the processes such as response to abiotic and biotic stimulus, cell organizations and biogenesis, and DNA or RNA metabolism (Figure 8). Liu et al. (2018) found the enrichment of heat-induced DMRs in the RNA metabolism and modification process, which may suggest the importance of RNA metabolism for stress adaptation. Zheng et al. (2017) reported that in rice, the long-term drought stress-induced epimutations were directly involved in the stress-responsive pathways. Wibowo et al. (2016) showed that hyperosmotic stress-induced DMRs were enriched with functions related to metabolic responses and ion transport; it was also suggested that exposure to hyperosmotic-stress targeted the discrete and epigenetically labile regions of the genome. The DMR-associated genes in the drought-tolerant introgression line DK151 were mainly involved in stress response, programmed cell death, and the nutrient reservoir activity, which may contribute to the constitutive drought tolerance (Wang et al., 2016). Stress priming can be utilised to develop epi-populations with epigenetic variations in various crops (Lämke and Bäurle 2017; Dalakouras and Vlachostergios 2021). Some of these stress associated epigenetic variations (DMPs/ DMRs) are stable and can faithfully transfer to the offspring (Zheng et al. 2017; Wibowo et al. 2016). These heritable stressed induced epigenetic variations epi-QTLs (DMPs and DMRs) can be used in epigenetic-assisted breeding and gene editing programs to develop superior next-generation stress tolerant crops (Dalakouras and Vlachostergios 2021; Gogolev et al 2021).

In conclusion, the stressed progeny derived from 25 generations of heat stress displayed a notable heat stress tolerance and higher genetic and epigenetic variations. Hence, our study suggests that the genetic and epigenetic plasticity induced by heat stress in the progeny might result in a phenotypic resilience to adverse environments and potentially trigger the process of microevolution under long-term stress conditions. Further study is required to discover the stable heritable genetic and epigenetic variations associated with heat stress resilience; a topic currently studied in our laboratory. The discovery of stress-associated stable heritable genetic and epigenetic markers could provide a potential source to develop a molecular breeding program for stress-tolerant crops.

## Acknowledgements

We thank Natural Sciences and Engineering Research Council of Canada (NSERC) for funding our work. Prof. Gideon Grafi from Ben-Gurion University of the Negev, Israel, has been acknowledged for scientific input on the manuscript draft.

## Author contributions

Narendra Singh Yadav– designed and performed experiments, analyzed data, wrote the manuscript

Viktor Titov– performed experiments and analyzed the data

Ivie Ayemere– performed experiments and analyzed the data

Boseon Byeon– conducted bioinformatics analysis

Yaroslav Ilnytskyy– conducted bioinformatics analysis

Igor Kovalchuk– came up with the concept of the study, analyzed data and contributed to writing.

## Materials & Correspondence

Correspondence and material requests should be addressed to corresponding author.

## Data availability

The sequencing data files from this study are deposited in the NCBI Sequence Read Archive (NCBI SRA) under BioProject - PRJNA802915 with accession number SRP357958. SRA identifier link: https://www.ncbi.nlm.nih.gov/bioprojet/?term=PRJNA802915

