## Supplementary material for "Multigenerational exposure to heat stress induces phenotypic resilience, and genetic and epigenetic variations in *Arabidopsis thaliana* offspring": Yadav et al 2022-Supplementary materials (from another version of this preprint: version 2, 2022-01-25)

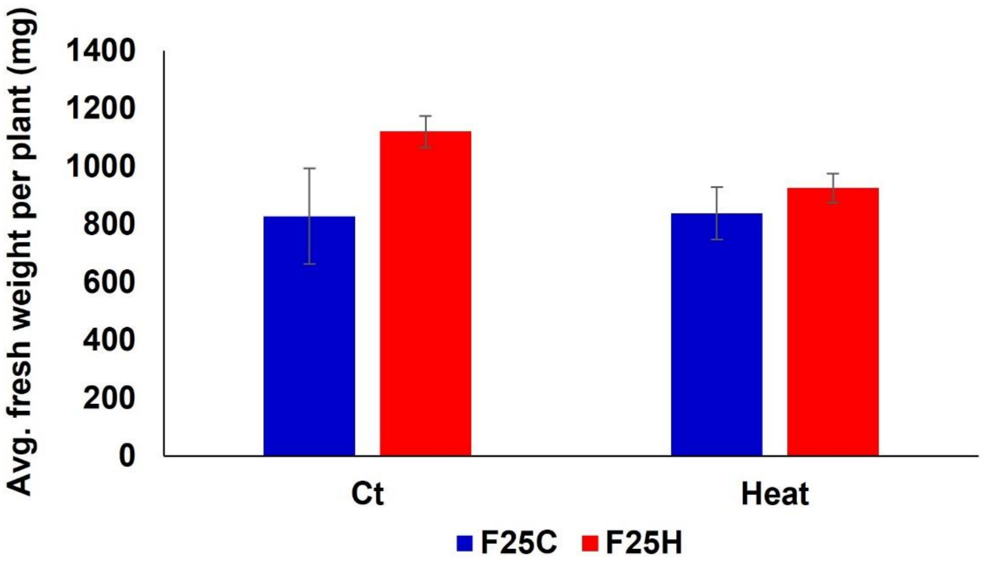


**Figure S1**. **Heat stress response at seedling stage**. The graph shows the average fresh weight (FW) in mg per plant (with ±SE). 'Ct' represents the plants grown under control condition and 'Heat' represents the plants grown under heat stress.

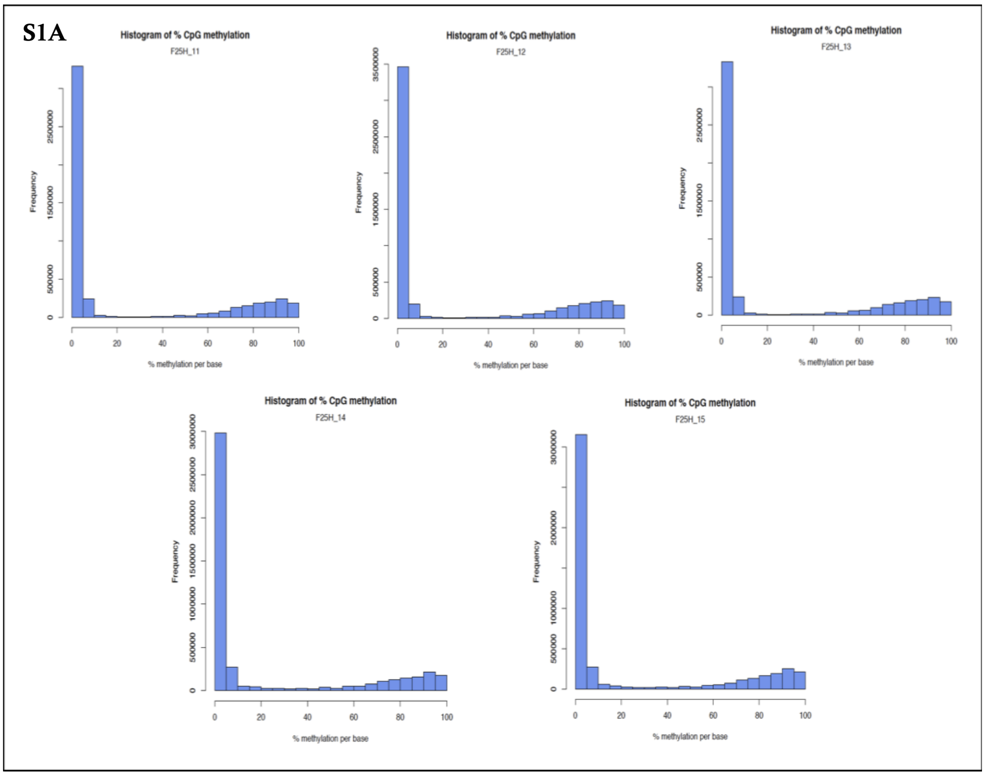
S1A
Histogram of % CpG methylation
F25H_11
Histogram of % CpG methylation
F25H_12
Histogram of % CpG methylation
F25H_13
Histogram of % CpG methylation
F25H_14
Histogram of % CpG methylation
F25H_15
Frequency
% methylation per base

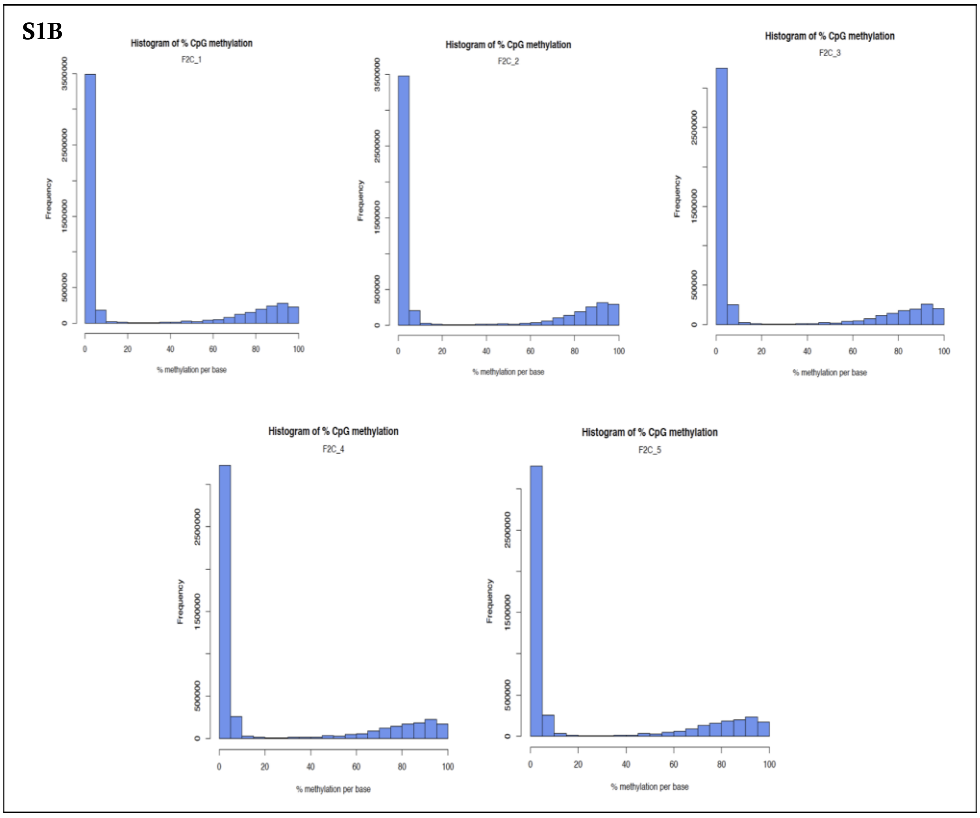
S1B
Histogram of % CpG methylation
F2C_1
Histogram of % CpG methylation
F2C_2
Histogram of % CpG methylation
F2C_3
Histogram of % CpG methylation
F2C_4
Histogram of % CpG methylation
F2C_5
Frequency
% methylation per base

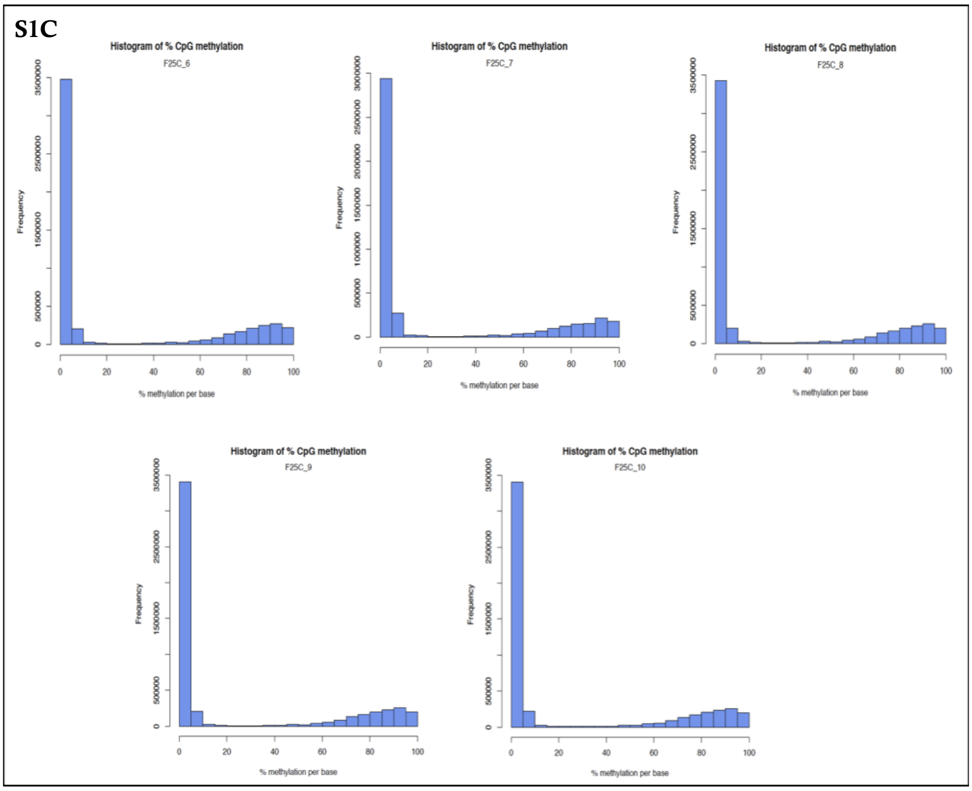


**Figure S2 A-C**. Frequency distribution of percentage methylation at CG sequence context for A) F25H, B) F2C, and C) F25C samples, most of the bases have either high or low methylation based on bimodal distribution. Data were normalized in *Methylkit* to account for clonal reads (PCR duplication bias), and each histogram represents a biological replicate, n=5 for F25H, F2C and F25H.

S2A

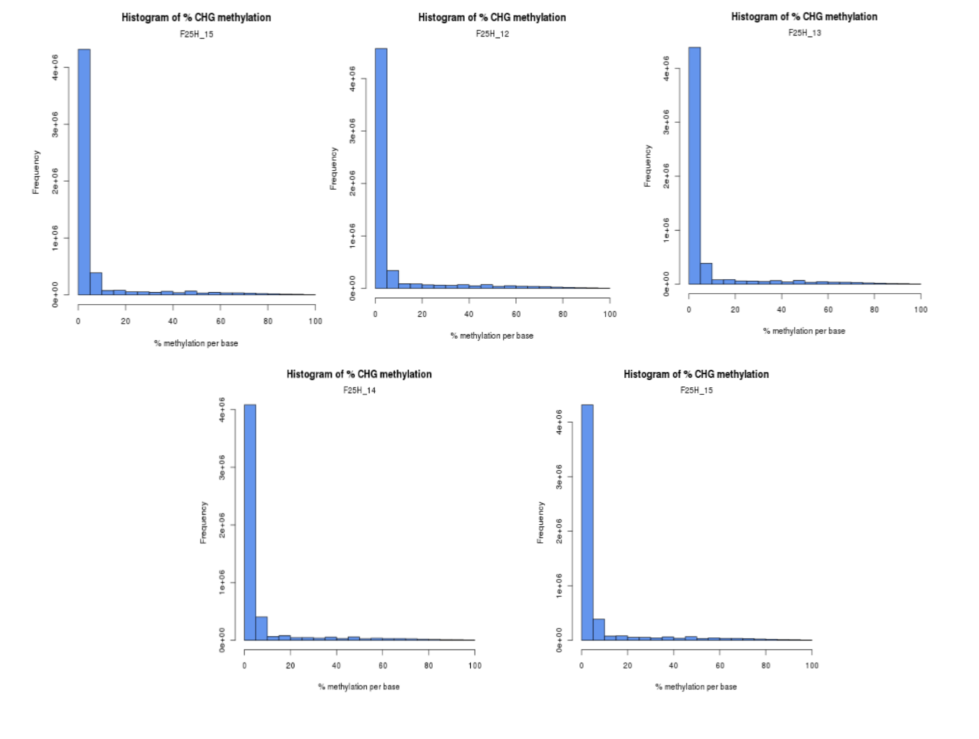
Histogram of % CHG methylation
F25H_15
Histogram of % CHG methylation
F25H_12
Histogram of % CHG methylation
F25H_13
Histogram of % CHG methylation
F25H_14
Histogram of % CHG methylation
F25H_15
Frequency
% methylation per base
0e+00
1e+06
2e+06
3e+06
4e+06
0
20
40
60
80
100

S2B

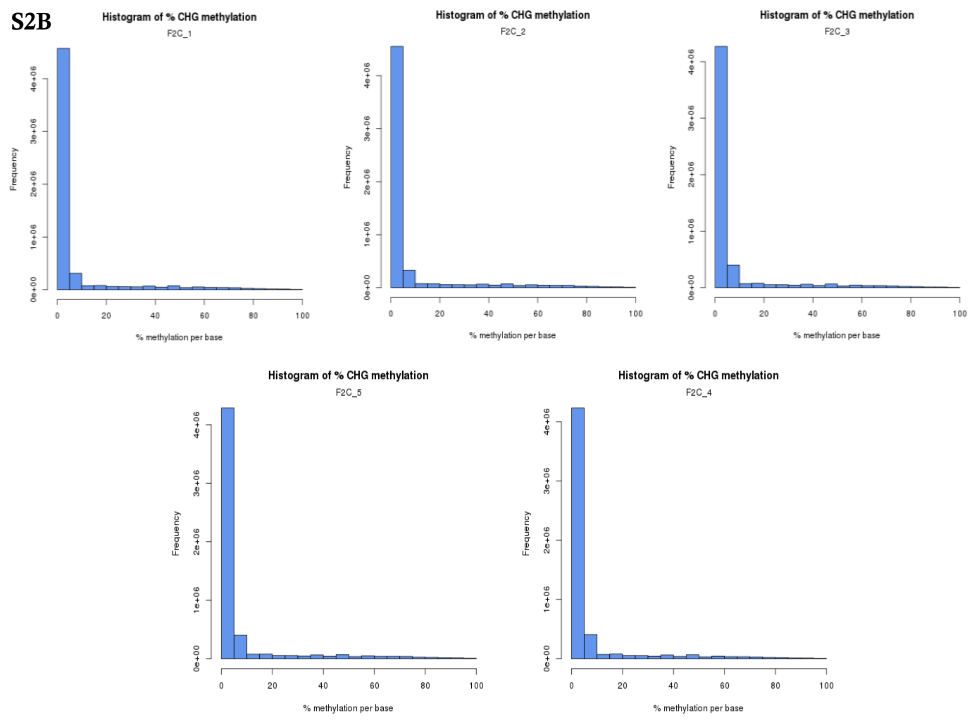
Histogram of % CHG methylation
F2C_1
Histogram of % CHG methylation
F2C_2
Histogram of % CHG methylation
F2C_3
Histogram of % CHG methylation
F2C_5
Histogram of % CHG methylation
F2C_4
Frequency
% methylation per base
0e+00
1e+06
2e+06
3e+06
4e+06
0
20
40
60
80
100

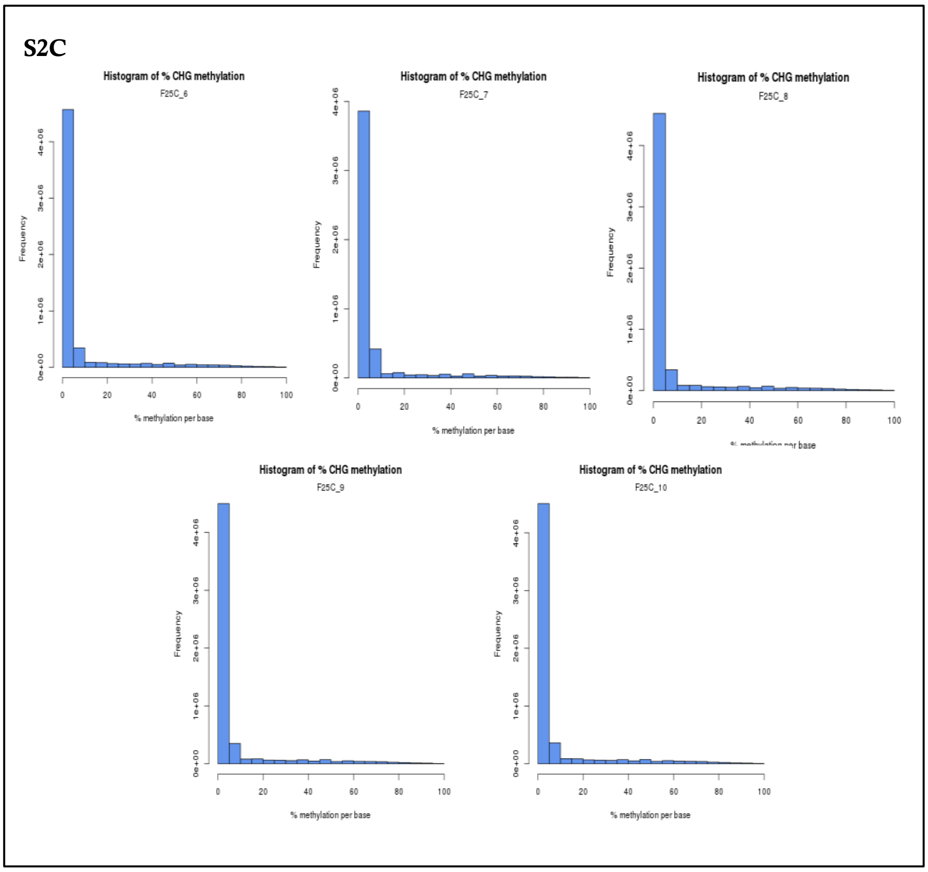


**Figure S3 A-C**. Frequency distribution of percentage methylation at CHG sequence context for A) F25H, B) F2C, and C) F25C samples, most of the bases have either high or low methylation based on bimodal distribution. Data were normalized in *Methylkit* to account for clonal reads (PCR duplication bias), and each histogram represents a biological replicate, n=5 for F25H, F2C and F25H. (H=A, T, C).

S3A

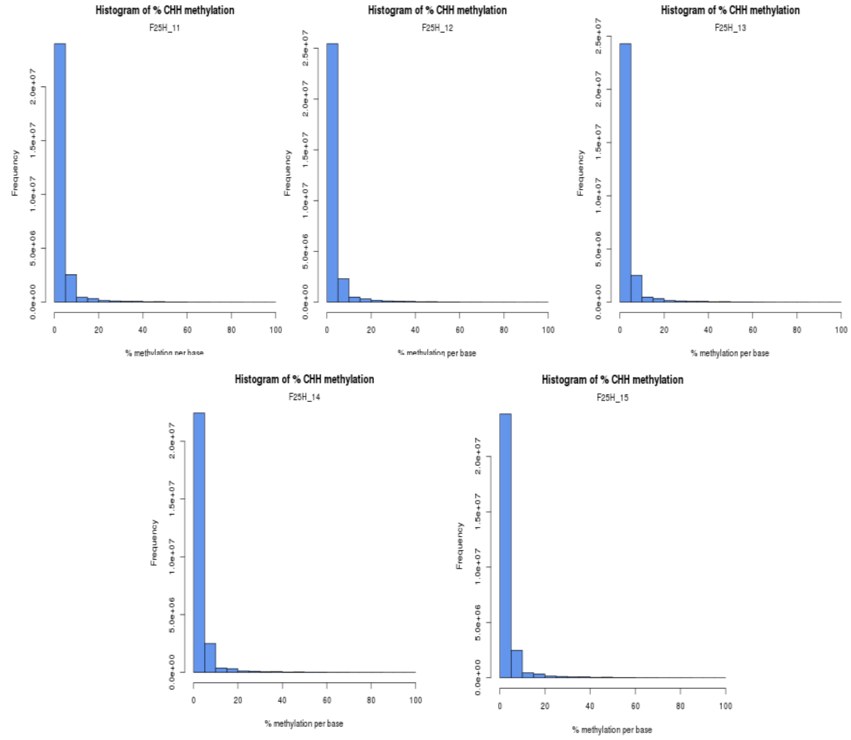
Histogram of % CHH methylation
F25H_11
Histogram of % CHH methylation
F25H_12
Histogram of % CHH methylation
F25H_13
Histogram of % CHH methylation
F25H_14
Histogram of % CHH methylation
F25H_15
Frequency
% methylation per base

S3B

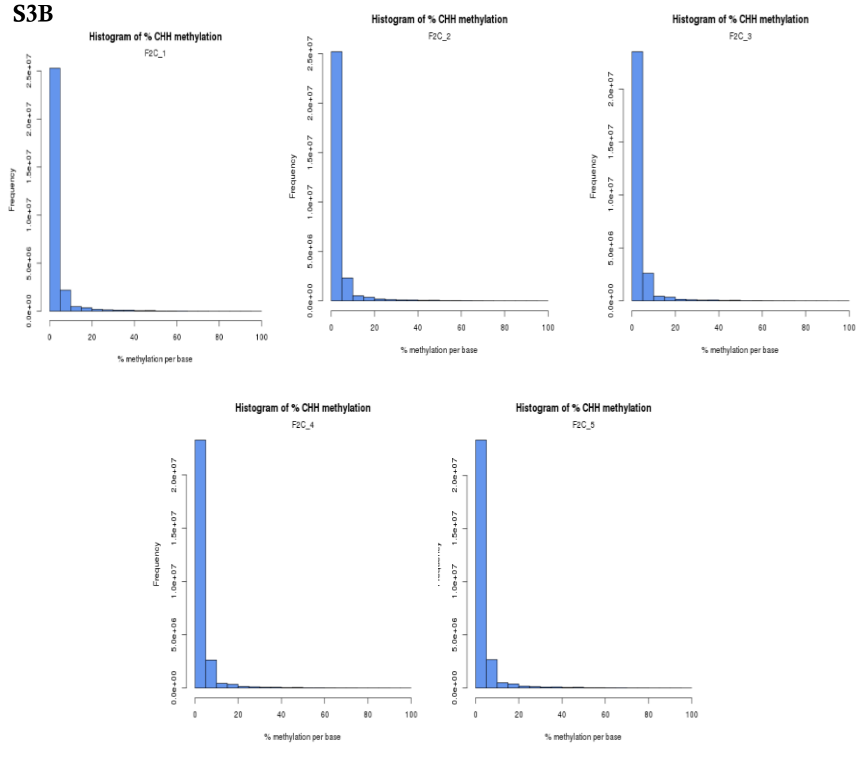
Histogram of % CHH methylation
F2C_1
Histogram of % CHH methylation
F2C_2
Histogram of % CHH methylation
F2C_3
Histogram of % CHH methylation
F2C_4
Histogram of % CHH methylation
F2C_5
Frequency
% methylation per base

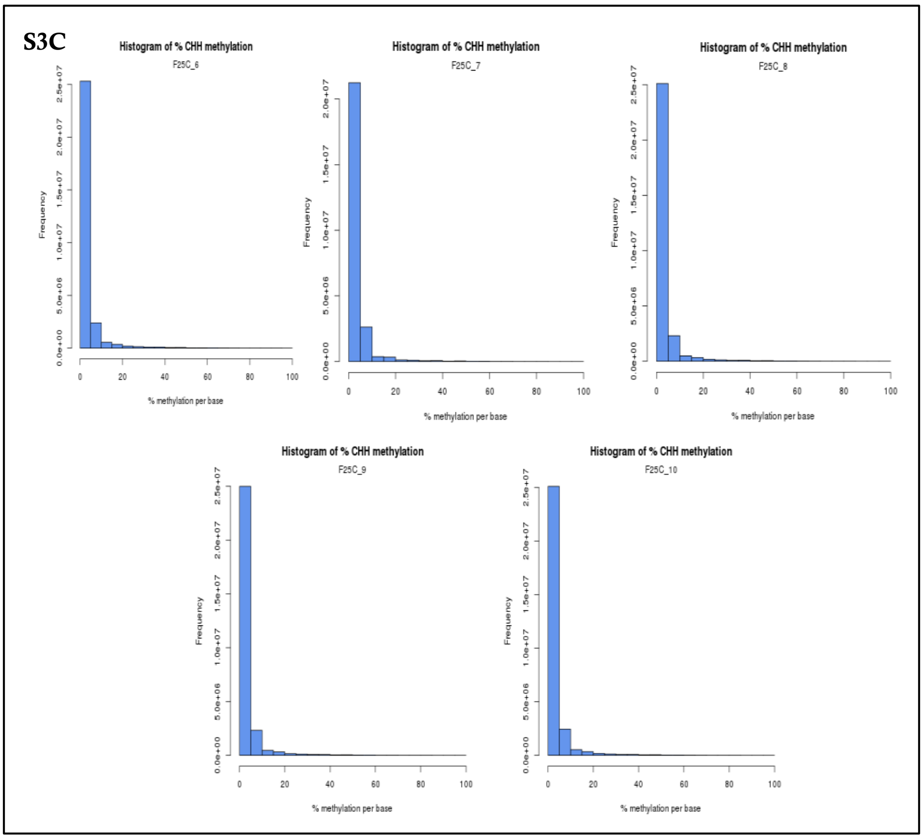


**Figure S4 A-C**. Frequency distribution of percentage methylation at CHH sequence context (H=A, T, C) for A) F25H, B) F2C, and C) F25C samples, most of the bases have either high or low methylation based on bimodal distribution. Data were normalized in *Methylkit* to account for clonal reads (PCR duplication bias), and each histogram represents a biological replicate, n=5 for F25H, F2C and F25H.

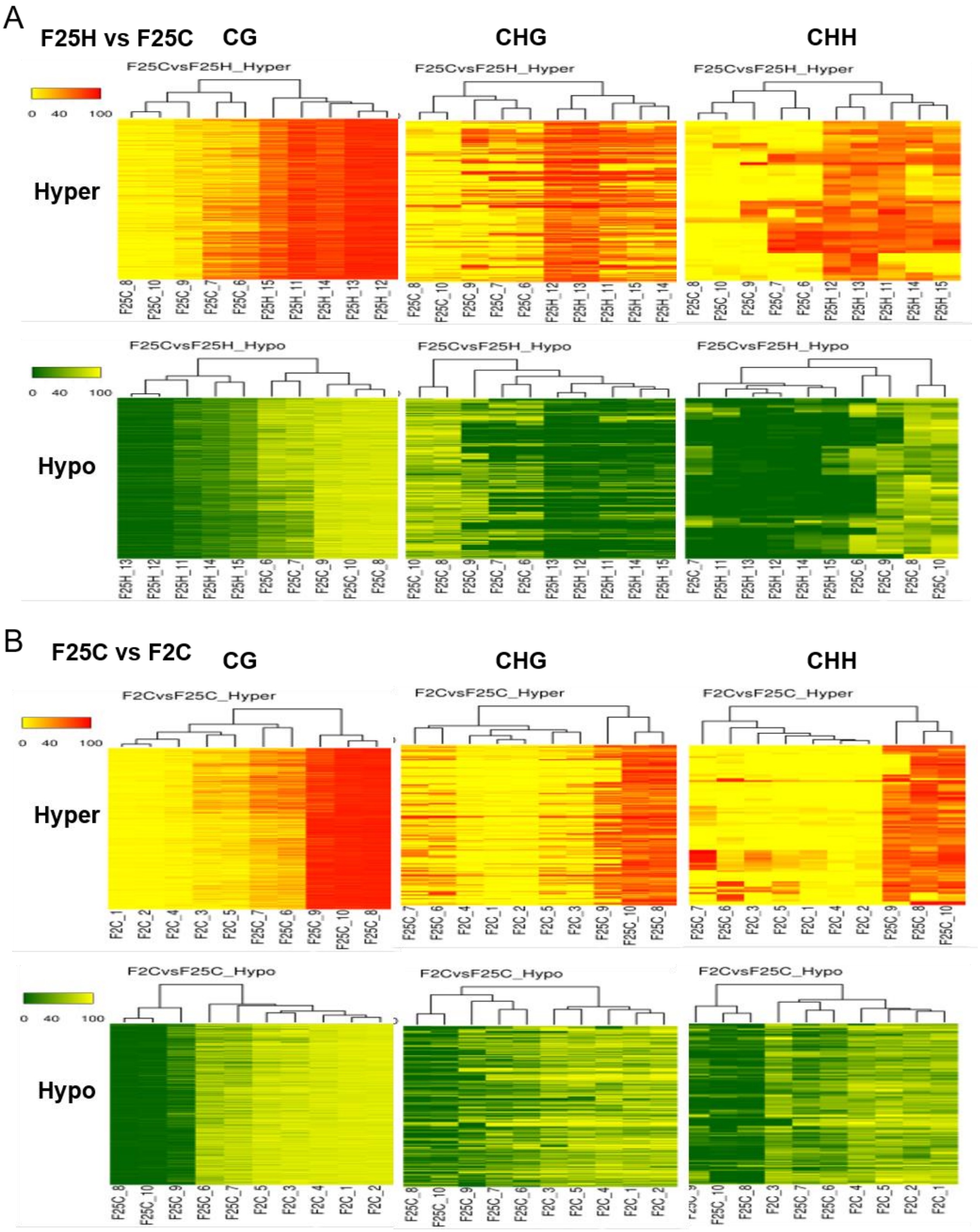


**Figure S5**. **A hierarchical clustering heatmap analysis.** (**A**) Heat maps of DMPs for hypermethylated cytosines (the upper panel) and hypomethylated cytosines (the lower panel) in CG, CHG and CHH contexts in F25H vs. F25C. Differentially methylated cytosines in the genome with differences > 50%

show the percentage of methylation in F25H vs. F2C. In ‘the upper panel’, the red section indicates the larger percentage of methylation, and the yellow section indicates the lower percentage, and in ‘the lower panel’, the green section indicates the larger percentage of methylation and the yellow one indicates the lower percentage, q-value <0.01. (**B**) Heat maps of DMPs for hypermethylated cytosines (the upper panel) and hypomethylated cytosines (the lower panel) in CG, CHG and CHH contexts in F25H vs. F2C. Differentially methylated cytosines in the genome with differences > 50% show the percentage of methylation in F25C vs. F2C. In ‘the upper panel’, the red section indicates the larger percentage of methylation, and the yellow section indicates the lower percentage, and in ‘the lower panel’, the green section indicates the larger percentage of methylation and the yellow one indicates the lower percentage, q-value <0.01.

***The distribution of DMPs and DMRs across the chromosomes***

The percentage of hypermethylated and hypomethylated DMPs per chromosome in the CG context showed an almost equal proportion of hyper- and hypo-methylation across all five chromosomes (Figure S5A). Hypermethylation and hypomethylation were apparent in the CG context (Figure S5A), while the CHG context (Figure S5A) had more hypermethylated bases distributed at the chromosome level in comparison groups of F25H vs F2C and F25C vs F2C. Hypomethylation was predominantly distributed in the CHH context (Figure S5A) and less distributed in the CHG context in all comparison groups. F25C vs. F2C had more hypermethylated bases than hypo-methylated bases per chromosome in the CHG context. In the CHH context, more hypomethylated bases per chromosome were found in all compared samples, except chromosome three which had an almost equal distribution of hyper- and hypomethylated bases (except F25H vs F2C) and the higher percentage of methylation than other chromosomes (Figure S5A). The proportions of DMPs per chromosome observed in F25H vs. F2C revealed more of hyper-methylated DMPs in the CHG context (Figure S5A), while in the CHH context, there were more hypo-methylated DMPs per chromosome (Figure S5A). The analysis of hyper/ hypo methylated events per chromosome indicates that F25H has a significant proportion ($q < 0.01$) of hyper-methylation in the CHG context (Figure S5A) and hypo-methylation in the CHH context (Figure S5A) compared with the non-stressed parental generation F2C. Similarly, the non-stressed parallel progeny F25C also shows a significant proportion ($q < 0.01$) of hyper-methylation compared with the parental generation F2C in the CHG context (Figure S5A) and hypo-methylation in the CHH context (Figure S5A).

The differentially extracted methylated regions (DMRs) show the same trends to differentially methylated positions (DMPs) in both the hyper- and hypomethylated contexts. The distribution of methylated regions highlights more hypermethylation in the CHG context and slightly more hypermethylation in the CG context; it also shows hypomethylation at the CHH context when F25H vs. F2C are compared in a 100 bp window (Figure S5B). When the sliding window was increased to 1000 bp, the proportion of hypermethylated DMRs per chromosome increased in CG and CHG contexts. DMRs in the CHH context in the 1000 bp sliding window showed no DMRs per chromosome (Figure S5C).

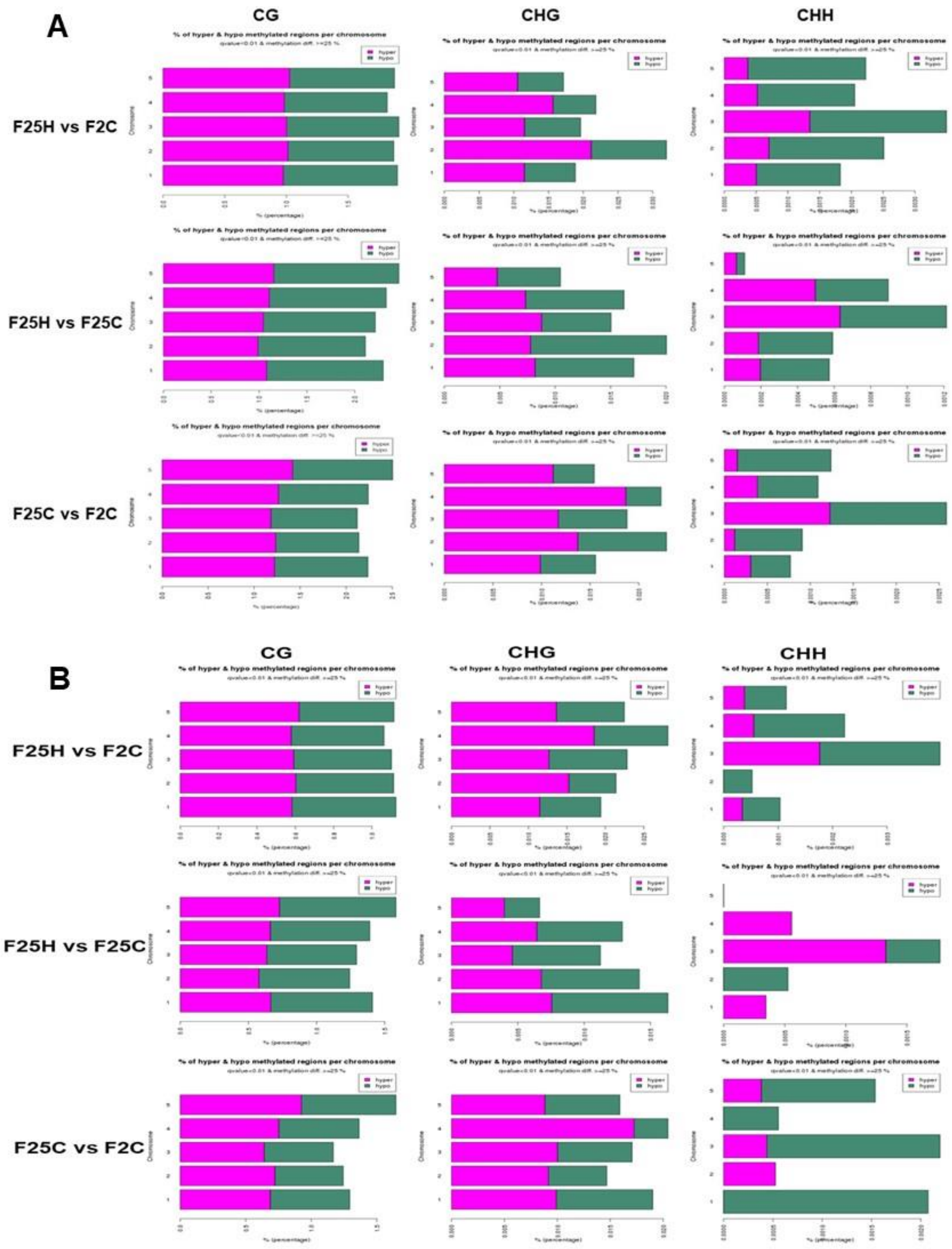
A
CG
CHG
CHH
F25H vs F2C
F25H vs F25C
F25C vs F2C
B
CG
CHG
CHH
F25H vs F2C
F25H vs F25C
F25C vs F2C

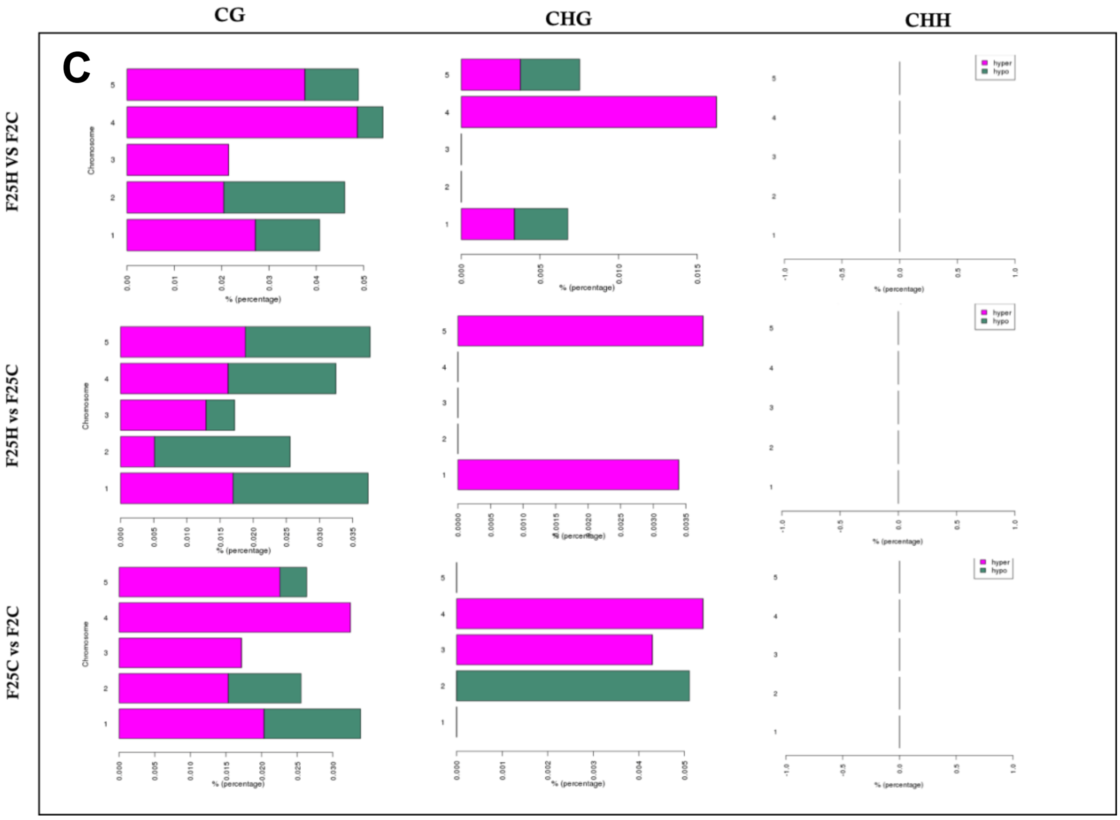


**Figure S6**. **The percentage of distributions of hyper- and hypo- methylation through the chromosome level**. (**A**) The distribution of DMPs in the CG context (the left panel), the CHG context (the middle panel) and the CHH context (the right panel) for each chromosome separately. The pink section indicates the percentage of hypermethylation, and the green one indicates hypomethylation, q-value <0.01 and the methylation difference > 25%. (**B**) The distribution of DMRs per chromosome per sequence context over a 100 bp window in the CG context (the left panel), in the CHG context (the middle panel) and in the CHH context (the right panel). DMRs with differences > 25% show the percentage of hyper- and hypomethylated regions; the pink section indicates the percentage of hypermethylation, and green sections indicates the percentage of hypomethylation, q-value <0.01. (**C**) Distribution of DMRs and percentage of hyper- and hypo- methylated bases at chromosome level DMRs per chromosome per sequence context over a 1000 bp window. Global DMRs with differences > 25% showing percentage of hyper- and hypomethylated regions pink section indicates the percentage of hypermethylation and green sections indicate the percentage of hypomethylation, q-value <0.01.

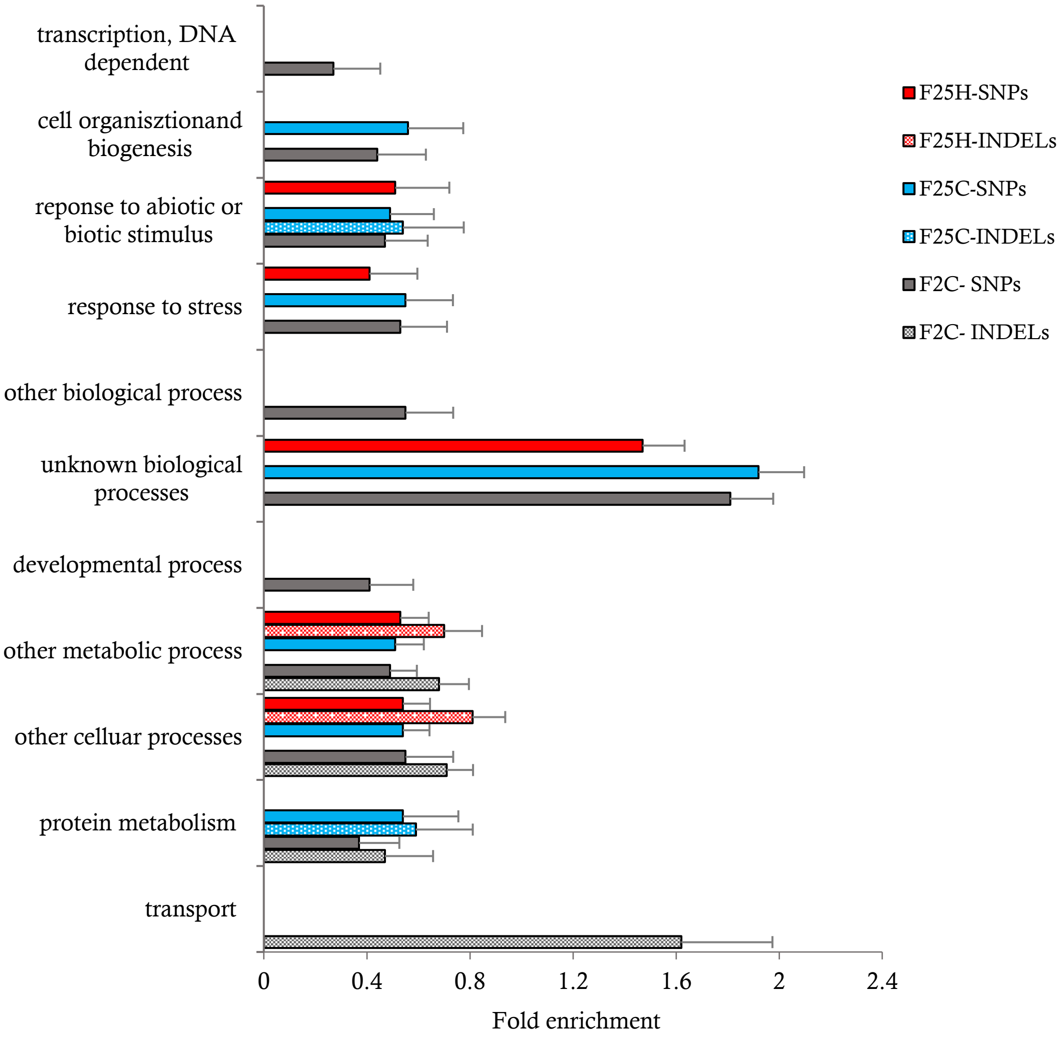


**Figure S7**. Enrichment analysis of SNPs and INDELs associated genes and their classification based on biological processes. y-axis is normalized Class Score with binomial coefficients as calculated by SuperViewer. Genes are classed with p-values < 0.05, ±bootstrap StdDev.

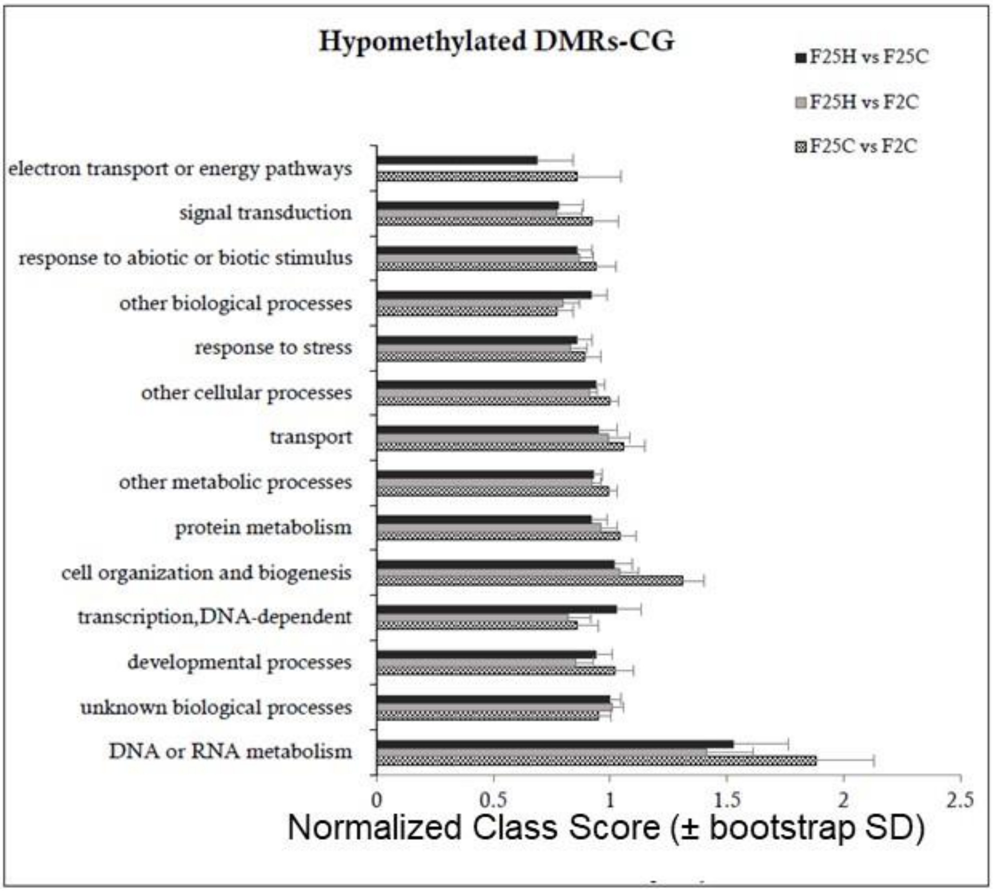


**Figure S8**. Enrichment analysis of hypomethylated DMRs on CG sites and their classification based on biological processes. Y-axis shows normalized Class Score with binomial coefficients as calculated by SuperViewer. To calculate enrichment, p-value of < 0.05, ±bootstrap StdDev were used.

A

B

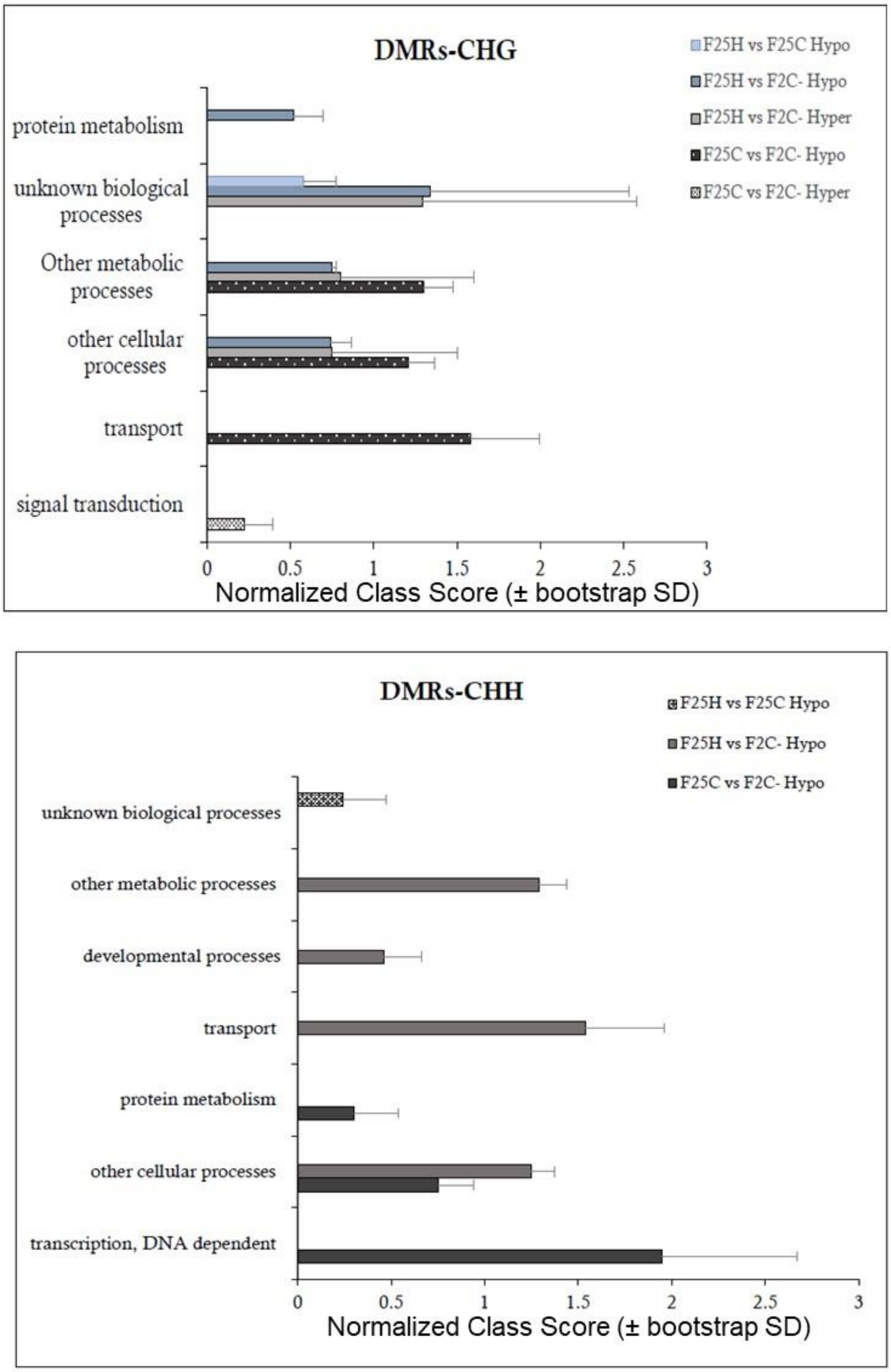


**Figure S9.** (A) Enrichment analysis of hypermethylated and hypomethylated DMRs on CHG sites and their classification based on biological processes. (B) Enrichment analysis of hypomethylated DMRs on CHG sites and their classification based on biological processes. Y-axis shows normalized Class Score with binomial coefficients as calculated by SuperViewer. To calculate enrichment, p-value of < 0.05, ±bootstrap StdDev were used.

**Table S1:** Total genetic variations (SNPs and INDELs) for F25H, F2C and F2C.

### Genetic variation analysis with relaxed parameters

#### Total genetic variations

| | SNPs | INDELs | Total |
|---|---|---|---|
| F25H | 53,678 | 17,669 | 71,347 |
| F2C | 15,599 | 7,986 | 23,585 |
| F25C | 15,526 | 8,143 | 23,669 |

#### Unique genetic variations

| | SNPs | INDELs | Total |
|---|---|---|---|
| F25H | 39,712 | 11,478 | 51,160 |
| F2C | 1,313 | 1,576 | 2,889 |
| F25C | 1,308 | 1,801 | 3,109 |

### Genetic variation analysis with strict parameters

#### Total genetic variations

| | SNPs | INDELs | Total |
|---|---|---|---|
| F25H | 12,000 | 6,177 | 18,177 |
| F2C | 5,190 | 4,643 | 9,833 |
| F25C | 5,188 | 4,781 | 9,969 |

#### Unique genetic variations

| | SNPs | INDELs | Total |
|---|---|---|---|
| F25H | 7,474 | 2,467 | 9,941 |
| F2C | 613 | 794 | 1,407 |
| F25C | 629 | 953 | 1,582 |

**Table S2: Classification of annotated SNPs and INDELs**

| Biological Processes | | F25H-SNPs | F25C – SNPs | F2C-SNPs | F25H-INDELs | F25C - INDELs | F2C-INDELs |
|---|---|---|---|---|---|---|---|
| Transport | Freq.<br>±SD<br>p-value | - | - | - | - | - | 1.62<br>±0.345<br>0.013 |
| Protein Metabolism | Freq.<br>±SD<br>p-value | - | 0.54<br>±0.215<br>0.000 | 0.37<br>±0.156<br>0.000 | - | 0.59<br>±0.221<br>0.000 | 0.47<br>±0.187<br>0.008 |
| Other Cellular Processes | Freq.<br>±SD<br>p-value | 0.54<br>±0.105<br>0.000 | 0.54<br>±0.103<br>0.000 | 0.55<br>±0.185<br>0.000 | 0.81<br>±0.127<br>0.043 | - | 0.71<br>±0.102<br>0.004 |
| Other Metabolic Processes | Freq.<br>±SD<br>p-value | 0.53<br>±0.11<br>0.000 | 0.51<br>±0.111<br>0.000 | 0.49<br>±0.104<br>0.000 | 0.7<br>±0.147<br>0.017 | - | 0.68<br>±0.116<br>0.004 |
| Developmental processes | Freq.<br>±SD<br>p-value | - | - | 0.41<br>±0.17<br>0.01 | - | - | - |
| Unknown biological process | Freq.<br>±SD<br>p-value | 1.47<br>±0.163<br>0.002 | - | 1.81<br>±0.167<br>0.000 | - | - | - |
| Other biological Processes | Freq.<br>±SD<br>p-value | - | 1.92<br>±0.177<br>0.000 | 0.55<br>±0.185<br>0.03 | - | - | - |
| Response to stress | Freq.<br>±SD<br>p-value | 0.41<br>±0.186<br>0.009 | - | 0.53<br>±0.181<br>0.02 | - | - | - |
| Response to abiotic or biotic stimulus | Freq.<br>±SD<br>p-value | 0.51<br>±0.21<br>0.026 | 0.55<br>±0.184<br>0.03 | 0.47<br>±0.166<br>0.02 | - | 0.54<br>±0.236<br>0.000 | - |
| Cell organization | Freq.<br>±SD<br>p-value | - | 0.49<br>±0.17<br>0.021 | 0.44<br>±0.189<br>0.02 | - | - | - |
| Transcription, DNA dependent | Freq.<br>±SD<br>p-value | - | 0.56<br>±0.214<br>0.043 | 0.27<br>±0.182<br>0.02 | - | - | - |

**Table S3: *Hypermethylated* DMRs enriched on CG Context**

| | | F25C vs F2C | F25H vs F2C | F25H vs F25C |
|---|---|---|---|---|
| DNA or *RNA* Metabolism | Freq.<br>±SD<br>p-value | 1.3<br>±0.203<br>0.009 | *1.43*<br>*0.018*<br>*0.002* | *1.65*<br>±0.201<br>0.000 |
| Unknown Biological Processes | Freq.<br>±SD<br>p-value | 1.05<br>±0.044<br>0.008 | *1.12*<br>±0.041<br>0.000 | *1*<br>±0.044<br>0.026 |
| Developmental Processes | Freq.<br>±SD<br>p-value | 1.02<br>±0.072<br>0.031 | *0.88*<br>±0.061<br>0.008 | *1.09*<br>±0.081<br>0.015 |
| *Transcription,DNA-dependent* | Freq.<br>±SD<br>p-value | 1<br>±0.098<br>0.041 | *0.85*<br>±0.087<br>0.014 | *0.88*<br>*±0.103*<br>*0.024* |
| Cell Organization and Biogenesis | Freq.<br>±SD<br>p-value | 0.89<br>±0.065<br>0.011 | *0.89*<br>±0.076<br>0.076 | *1.23*<br>±0.087<br>0.001 |
| Protein Metabolism | Freq.<br>±SD<br>p-value | 0.89<br>±0.066<br>0.007 | *0.84*<br>*±0.067*<br>*0.002* | *1.02*<br>*±0.075*<br>*0.031* |
| Other Metabolic Processes | Freq.<br>±SD<br>p-value | 0.88<br>±0.034<br>0.000 | *0.85*<br>±0.038<br>0.000 | *0.96*<br>±0.036<br>0.016 |
| Transport | Freq.<br>±SD<br>p-value | 0.88<br>*±0.072*<br>*0.012* | *0.93*<br>±0.078<br>0.028 | *1.08*<br>±0.078<br>0.023 |
| Other Cellular Processes | Freq.<br>±SD<br>p-value | 0.87<br>±0.031<br>0.000 | *0.85*<br>±0.031<br>0.000 | *0.96*<br>±0.036<br>0.015 |
| Response to Stress | Freq.<br>±SD<br>p-value | 0.83<br>±0.055<br>0.001 | *0.89*<br>±0.071<br>0.009 | *1*<br>±0.07<br>0.034 |
| Other Biological Processes | Freq.<br>±SD<br>p-value | 0.82<br>±0.066<br>0.001 | *0.77*<br>*±0.067*<br>*0.000* | *0.86*<br>±0.068<br>0.006 |
| Response to Abiotic or Biotic Stimulus | Freq.<br>±SD<br>p-value | 0.78<br>±0.057<br>0.000 | *0.8*<br>±0.071<br>0.000 | *1.03*<br>±0.08<br>0.032 |
| Signal Transduction | Freq.<br>±SD<br>p-value | 0.73<br>±0.095<br>0.001 | *0.69*<br>±0.11<br>0.001 | *0.84*<br>±0.106<br>0.022 |
| Electron Transport or Energy Pathways | Freq.<br>±SD<br>p-value | 0.87<br>±0.133<br>0.004 | *0.56*<br>±0.144<br>0.003 | *0.51*<br>±0.129<br>0.002 |

**Table S4: *Hypomethylated* DMRs enriched on CG Context**

| | | F25C vs F2C | F25H vs F2C | F25H vs F25C |
|---|---|---|---|---|
| DNA or RNA Metabolism | Freq.<br>±SD<br>p-value | 1.88<br>±0.25<br>0.000 | 1.41<br>±0.201<br>0.006 | 1.53<br>±0.234<br>0.000 |
| Unknown Biological Processes | Freq.<br>±SD<br>p-value | 0.95<br>±0.053<br>0.016 | 1.01<br>±0.047<br>0.027 | 1<br>±0.044<br>0.025 |
| Developmental Processes | Freq.<br>±SD<br>p-value | 1.02<br>±0.081<br>0.035 | 0.85<br>±0.077<br>0.006 | 0.94<br>±0.071<br>0.026 |
| Transcription, DNA-dependent | Freq.<br>±SD<br>p-value | 0.86<br>±0.087<br>0.02 | 0.82<br>±0.095<br>0.012 | 1.03<br>±0.102<br>0.039 |
| Cell Organization and Biogenesis | Freq.<br>±SD<br>p-value | 1.31<br>±0.091<br>0.000 | 1.04<br>±0.084<br>0.034 | 1.02<br>±0.076<br>0.033 |
| Protein Metabolism | Freq.<br>±SD<br>p-value | 1.04<br>±0.069<br>0.027 | 0.96<br>±0.072<br>0.032 | 0.92<br>±0.067<br>0.018 |
| Other Metabolic Processes | Freq.<br>±SD<br>p-value | 0.99<br>±0.038<br>0.025 | 0.92<br>±0.04<br>0.005 | 0.93<br>±0.036<br>0.004 |
| Transport | Freq.<br>±SD<br>p-value | 1.06<br>±0.089<br>0.029 | 0.99<br>±0.096<br>0.041 | 0.95<br>±0.081<br>0.033 |
| Other Cellular Processes | Freq.<br>±SD<br>p-value | 1<br>±0.035<br>0.025 | 0.91<br>±0.035<br>0.002 | 0.94<br>±0.034<br>0.005 |
| Response to Stress | Freq.<br>±SD<br>p-value | 0.89<br>±0.072<br>0.012 | 0.83<br>±0.073<br>0.003 | 0.86<br>±0.063<br>0.004 |
| Other Biological Processes | Freq.<br>±SD<br>p-value | 0.77<br>±0.072<br>0.00 | 0.8<br>±0.068<br>0.001 | 0.92<br>±0.068<br>0.02 |
| Response to Abiotic or Biotic Stimulus | Freq.<br>±SD<br>p-value | 0.94<br>±0.083<br>0.027 | 0.87<br>±0.056<br>0.009 | 0.86<br>±0.064<br>0.005 |
| Signal Transduction | Freq.<br>±SD<br>p-value | 0.92<br>±0.118<br>0.046 | 0.77<br>±0.108<br>0.011 | 0.78<br>±0.106<br>0.007 |
| Electron Transport or Energy Pathways | Freq.<br>±SD<br>p-value | 0.86<br>±0.184<br>0.000 | - | 0.69<br>±0.151<br>0.02 |

**Table S5: DMRs enriched on CHG Context**

| | | Hypomethylated | | | Hypermethylated | |
|---|---|---|---|---|---|---|
| | | F25C<br>vs<br>F2C | F25H<br>vs<br>F2C | F25H<br>vs<br>F25C | F25C<br>vs<br>F2C | F25H<br>vs<br>F2C |
| Signal Transduction | Freq.<br>±SD<br>p-value | - | - | - | 0.22<br>±0.173<br>0.048 | - |
| Transport | Freq.<br>±SD<br>p-value | 1.58<br>±0.411<br>0.41 | - | - | - | - |
| Other Cellular Processes | Freq.<br>±SD<br>p-value | 1.21<br>±0.164<br>0.04 | 0.74<br>±0.015<br>0.043 | - | - | 0.75<br>±0.146<br>0.021 |
| Other Metabolic Processes | Freq.<br>±SD<br>p-value | 1.3<br>±0.179<br>0.02 | 0.75<br>±0.024<br>0.024 | - | | 0.8<br>±0.164<br>0.043 |
| Unknown Biological Processes | Freq.<br>±SD<br>p-value | - | 1.34<br>±1.191<br>0.015 | 0.58<br>±0.196<br>0.02 | - | 1.29<br>±0.208<br>0.027 |
| Protein Metabolism | Freq.<br>±SD<br>p-value | - | 0.52<br>±0.175<br>0.029 | - | - | - |

**Table S6: Hypomethylated DMRs Enriched on CHH Context**

| | | F25C vs F2C | F25H vs F2C | F25H vs F25C |
|---|---|---|---|---|
| Transcription, DNA Dependent | Freq. ±SD p-value | 1.95 ±0.721 0.04 | - | - |
| Other Cellular Processes | Freq. ±SD p-value | 0.75 ±0.189 0.049 | 1.25 ±0.126 0.01 | - |
| Protein Metabolism | Freq. ±SD p-value | 0.3 ±0.235 0.24 | - | - |
| Transport | Freq. ±SD p-value | - | 1.54 ±0.417 0.03 | - |
| Developmental Processes | Freq. ±SD p-value | - | 0.46 ±0.2 0.02 | - |
| Other Metabolic Processes | Freq. ±SD p-value | - | 1.29 ±0.146 0.01 | - |
| Unknown Biological Processes | Freq. ±SD p-value | - | - | 0.24 ±0.232 0.05 |
